# Alpha activity neuromodulation induced by individual alpha-based neurofeedback learning in ecological context: A double-blind randomized study

**DOI:** 10.1101/2020.12.02.406173

**Authors:** Fanny Grosselin, Audrey Breton, Lydia Yahia-Cherif, Xi Wang, Giuseppe Spinelli, Laurent Hugueville, Philippe Fossati, Yohan Attal, Xavier Navarro-Sune, Mario Chavez, Nathalie George

## Abstract

The neuromodulation induced by neurofeedback training (NFT) remains a matter of debate. Investigating the modulation of brain activity specifically associated with NF requires controlling for multiple factors, such as reward, performance, congruency between task and targeted brain activity. This can be achieved using sham feedback (FB) control condition, equating all aspects of the experiment but the link between brain activity and FB. We aimed at investigating the modulation of individual alpha EEG activity induced by NFT in a double-blind, randomized, sham-controlled study. Forty-eight healthy participants were assigned to either NF (n=25) or control (n=23) group and performed alpha upregulation training (over 12 weeks) with a wearable EEG device. Participants of the NF group received FB based on their individual alpha activity. The control group received the auditory FB of participants of the NF group. An increase of alpha activity across training sessions was observed in the NF group only (*p<0.001*). This neuromodulation was selective in that it was not observed for theta (4-8Hz) and low beta (13-18Hz) activities. While alpha upregulation was found in the NF group only, psychological outcome variables showed increased feeling of control, decreased anxiety level and increased relaxation feeling in both the NF and control groups, without any significant difference between groups. This is interpreted in terms of learning context and placebo effects. Our results pave the way to self-learnt, NF-based neuromodulation with light-weighted, wearable EEG systems.

## Introduction

Neurofeedback (NF) is a cognitive training that exploits the causal relationship between brain activity and cognitive-motor abilities. As brain-computer interfaces (BCI) applications, NF consists in providing real-time auditory, visual, or tactile feedback of a subject’s brain activity to train self-regulation of specific brain patterns related to a targeted ability. NF applications have been developed since the 70’s in non-clinical ^1–3^ and clinical settings, such as epilepsy ^4^, attention-deficit hyperactivity disorder ^5–9^, depression ^10,11^, psychopathy ^12,13^, and anxiety ^10,14–16^. However, the neurocognitive mechanisms underlying BCI tasks or NF training (NFT) remain elusive ^17,18^. The neuromodulation associated with NFT has already been studied in several contexts ^19–21^, but this was not yet done in a long-term, multiple-session (12 weeks), sham-controlled design using an ecological reinforcer NF context for both NF and control groups.

In the previous literature, control conditions are quite variable in NF studies, not only aiming at the link between brain activity and feedback but also varying the task or the procedure ^22^. For example, in clinical studies where NFT aimed at reducing behavioral symptoms or psychological processes associated with various disorders (anxiety ^14–16^, depression ^10,11,23^, addiction ^24–27^, attention deficit ^5–9,28,29^), NF performance was typically compared with active control groups, such as cognitive therapy, mental exercise, and treatment-as-usual ^22^. Thus, the self-reported or clinical benefits of NFT may be related to an ensemble of specific and non-specific mechanisms, including psychosocial influences ^30–32^, cognitive and attentional/motivational factors ^33^, test-retest improvement, as well as spontaneous clinical improvement or cognitive development ^17^, and the learning context, contributing to the ongoing debate about NF efficacy ^15,34^. In some NF studies, the control condition was based on linking the feedback to another brain activity than the targeted one ^22,35^ which entails an incongruity between the activity driving the feedback and the task—hence the cognitive efforts—of the subject. Here, we used a sham feedback (sham-FB) condition for the control group—as it is commonly used in other studies including MEG or fMRI NF protocols ^19–22,36–39^. The participants in the sham-FB group received ‘yoked’ feedback, corresponding to the feedback of randomly-chosen subjects from the NF group at the same stage of learning. Hence, this feedback was similar in every aspect to the one in the NF group, except that it was not the result of an established link between the subject’s alpha-band activity and the auditory stream. Such sham-FB control condition breaks the operant link between the subject’s neuromodulation and the received feedback, which may be seen as its main limitation ^22,40^. Yet, this operant link is considered as constitutive of NFT and its effects ^41^, and this sham-FB control condition has the advantage to allow matching for reward and performance across the control and the NF groups ^22^. Thus, it allows controlling as closely as possible for the learning context while breaking the operant learning component that is key to NFT.

Some studies already used alpha up-regulation NFT for improving different cognitive processes such as episodic memory ^36^ or mental performances ^42^. Moreover, across-sessions neuromodulation making use of sham-controlled design was tested in the past—not necessarily targeting alpha. However, some of these studies included only one or a few sessions ^37,39^ and others did not find clear evidence of across-sessions neuromodulation ^19,21,38^. In addition, in these studies, the learning context and task were not directly linked to the expected cognitive performance or targeted psychological process ^21,36^. In the present study, we used an ‘ecological’ context with respect to the NF task, in which all participants were asked to close their eyes and get immersed in a relaxing soundscape delivered by headphones, while being engaged in the task. This task consisted in learning to decrease the volume of a sound indicator, which was inversely related to their individual alpha-band EEG activity recorded by two parietal dry electrodes. This conditioned all the participants of the two groups to relax and increase their alpha EEG band activity—as alpha activity is known to be linked with resting, relaxed or meditative states ^43–50^. This constituted a ‘transparent’ context ^18^, which may be essential to unravel the mechanisms of NF learning. This matched learning context between the NF and control group allowed us to rigorously test the alpha-band neuromodulation specifically induced by NF.

Most of the NF studies are performed using wet EEG sensors in a laboratory context. Here, we used a new compact and wearable EEG-based NF device with dry electrodes (Melomind™, myBrain Technologies, Paris, France). This device was studied in an under-review study by a comparison to a standard-wet EEG system (Acticap, BrainProducts, Gilching, Germany) ^51^. As suggested in ^52–54^, novel low-cost dry electrodes have comparable performances in terms of signal transfer for BCI and can be suitable for EEG studies. Moreover, such a user-friendly and affordable device with few dry sensors, does not require conductive gel, and can be so suitable for easy real-life use by the general population.

Here, we aimed at studying the neuromodulation specifically induced by individual alpha up-regulation NFT over multiple sessions throughout 12 weeks, in a double-blind, randomized, sham-controlled design study within general healthy population in an ecological reinforcement context. As in many NF protocols aiming at anxiety reduction, stress-management or well-being ^14,15,55,56^, we chose an alpha-upregulation NFT for the known link between the increase of low frequency EEG activities—including theta and alpha activities—and relaxed or meditative states ^43–49^.

We expected an increase of the trained individual alpha band activity across sessions in the NF group, because it was asked to the participants to find their own strategies to reduce the volume of the auditory feedback—operantly linked to the individual alpha activity in the NF group only—and this, as a learning process, requires multiple sessions ^41,57^. The use of an ecological, relaxing, learning context, allowed us to test if alpha upregulation could be induced just by the context, in which case, we should observe an increase of alpha activity in both the NF and control groups. In contrast, if alpha neuromodulation is specific to the NFT, one could expect a significant increase of alpha activity across sessions only in the NF group. Finally, we were also interested in the impact of such NFT on self-reports related to anxiety level and relaxation. The improvement of such self-reported psychological processes can be due to specific NF mechanisms and non-specific mechanisms ^17^, such as the context of the learning including instructions, the biomarker ^58^ used and psychosocial factors^33^. Considering the learning context (relaxing auditory landscape) and instructions (closed eyes during 21 minutes) that we used in both groups, associated with the sham-controlled design in a healthy population, we expected improvements in relaxation and anxiety in the NF group and in the control group due to placebo effects ^30,59^.

## Materials and Methods

### Participants

In the NF literature, the common number of included subjects varies from 10 to 20 participants by group ^21,35–37,60^. Based on the literature and resources constraints ^61^ implying a follow-up across 12 weeks for each subject (see the “Experimental protocol” section), forty-eight healthy volunteers, divided in two groups, were included in this study (mean age: 33.3 years; age range: 18-60; see Supplementary Table S1 for more details). All participants declared having normal or corrected-to-normal vision, no hearing problem, no history of neurological or psychiatric disorders, no ongoing psychotropic drug treatment and no or little NF or BCI experience. Participants were blindly assigned either to the *NF group*—who received real NF—or to the *control group*—who received sham-FB. For the purpose of the sham-FB design construction, the first *N* participants were assigned to the *NF group*. Only the experimenters and the data analysts knew the existence of the two groups and that the first *N* subject(s) was/were in the *NF group*. However, the experimenters and the data analysts were blind to *N* and blind to the random assignment after *N*. This resulted in a double-blind sham-controlled design with 25 subjects in the *NF group* and 23 in the *control group*. The blind assignment was maintained until the end of the experiment. No test was done to know if the participants suspected the existence of two groups and their assignment to one of these groups.

Participants were enrolled from the general population through advertisements in science and medical schools in Paris, through an information mailing list (RISC, https://expesciences.risc.cnrs.fr) and through flyers distributed in companies in Paris (France).

Participants completed the protocol in three different locations: at the Center for NeuroImaging Research (CENIR) of the Paris Brain Institute (ICM, Paris, France) (N=20 participants, *NF group*: 10, *control group*: 10), at their workplace 14 (N=14, *NF group*: 8, *control group*: 6), or at home (N=14, *NF group*: 7, *control group*: 7). The 20 participants who performed the protocol at the CENIR were part of those planned in the study approved by French ethical committee (CPP Sud-Ouest et Outre Mer I, ref. 2017-A02786-47), registered on ClinicalTrials.gov (ref. NCT04545359), although the present study was not part of this clinical study. For these participants, a financial compensation was provided at the end of the study for the time taken to come to the lab. The 28 other participants followed the same protocol but performed it in a real-life context (at work, at home). Moreover, all participants gave written and informed consent in accordance with the Declaration of Helsinki.

### EEG recording and preprocessing

Brain activity was recorded by two gold-coated dry electrodes placed on parietal regions (P3 and P4) (Melomind™, myBrain Technologies, Paris, France; Fig. 1). Ground and reference were silver fabric electrodes, placed on the left and right headphones respectively, in mastoid regions.

**Figure 1.**
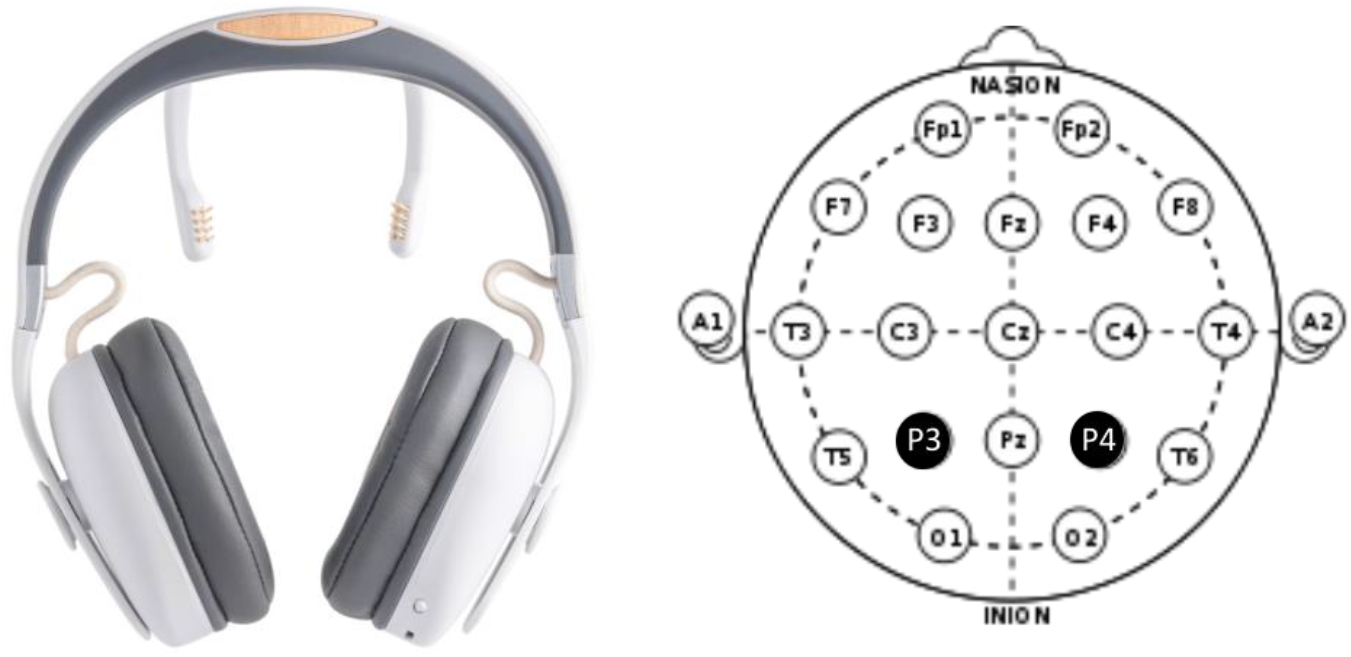
Melomind™ device. *Left*: The device comprises an audio headset and two posterior branches at the end of which two gold-coated dry electrodes are inserted. *Right*: These electrodes are positioned at P3 and P4 locations (indicated in black) according to the extended 10-20 International System.

EEG signals were amplified and digitized at a sampling rate of 250 Hz, band-pass filtered from 2 Hz to 30 Hz in real-time and sent to the mobile device by embedded electronics in the headset. The headset communicated via Bluetooth with a mobile NF application, which processed EEG data every second to give the user auditory feedback about his/her alpha-band activity (see below). A DC offset removal was applied on each second of data for each channel and a notch filter centered at 50Hz was applied to remove the powerline noise. Real-time estimation of signal quality was then performed by a dedicated machine learning algorithm ^62^. Briefly, this algorithm computed in time and frequency domains, EEG measures that are commonly used in artefact detection from electrophysiological signals (standard deviation, skewness, kurtosis, EEG powers in different frequency bands, power of change, etc.). These EEG features were compared to a training database by a k-nearest neighbors classifier to assign a quality label to the EEG signal among three classes: HIGHq, MEDq, and LOWq (see ^62^ for more details). In Grosselin et al., 2019 ^62^, we showed that this algorithm has an accuracy higher than 90% for all the studied databases. This algorithm was used to detect noisy segments (LOWq) which were excluded from posterior analysis.

### Experimental protocol

Based on previous studies ^15,63^, we proposed a protocol consisting in 12 NFT sessions, with one session per week (Fig. 2). Each session was composed of 7 exercises of 3 minutes (total: 21 minutes), which corresponded to 4.2 hours of training. At the beginning and end of each session, two-minute resting state recording was performed and the participant completed the Spielberger State-Trait Anxiety Inventory (*STAI*, *Y-A* form, in French, ^64^)—to assess his/her anxiety state level—and a 10-cm visual analog scale (VAS) indicating his/her subjective relaxation level (r*elax-VAS*). These resting state recordings were not analyzed here as they are out of the scope of this study focused on neuromodulation. Moreover, at the end of each 3-minute exercise, the participant indicated his/her subjective level of feedback control on a 10-cm VAS (*control-VAS*)—the left side indicating the feeling of no control; the right bound indicating a feeling of perfect control.

**Figure 2.**
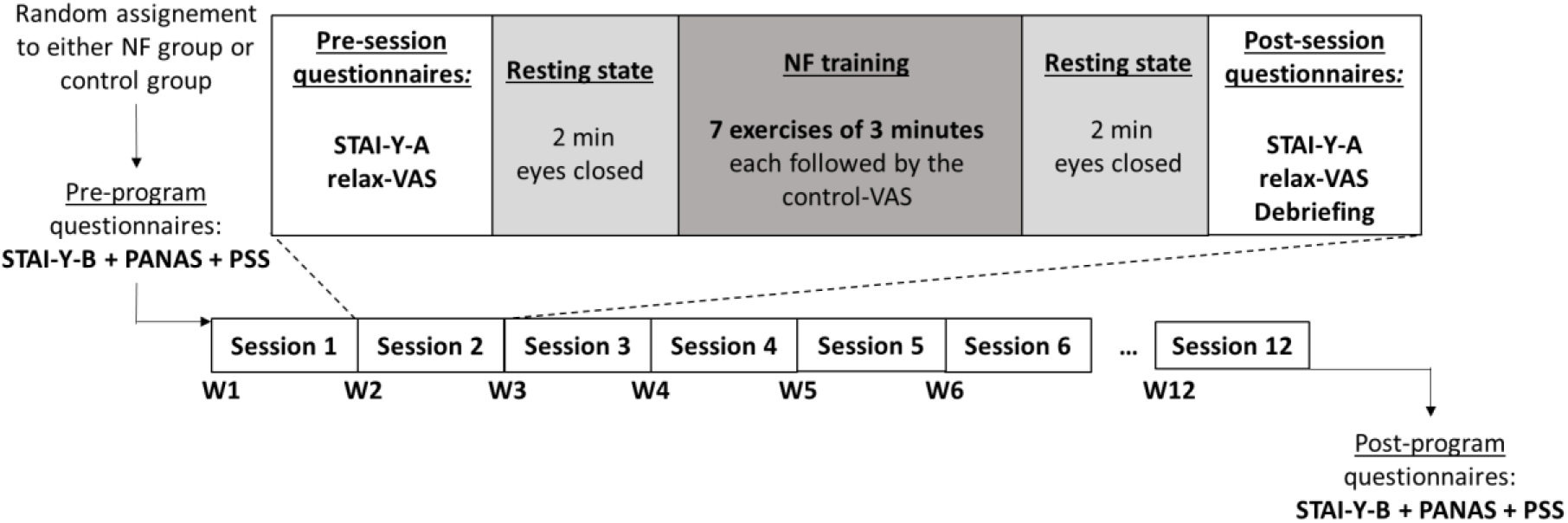
Timeline of the experimental protocol. Before and after the NFT program, the participants completed three psychometric trait questionnaires, to assess psychological stress (Perceived Stress Scale - PSS) ^65^, trait anxiety (*STAI-Y-B*) ^64^, and affectivity (Positive and Negative Affect Schedule - PANAS). NFT was performed over 12 weeks (W1, W2, … W12), with one training session per week (Session 1, …, Session 12). See main text for detailed description of the sessions.

### Neurofeedback training procedure

The NF paradigm targeted alpha rhythm centered on the individual alpha frequency (IAF). Before each NFT session, a 30-second calibration phase allowed computing IAF using an optimized, robust estimator dedicated to real-time IAF estimation based on spectral correction and prior IAF estimations ^66^. More precisely, the spectrum was corrected by removing the 1/f trend estimated by an iterative curve-fitting procedure. Then, local maxima were detected in the corrected spectrum between 6 and 13 Hz as the downward going zero crossing points in the first derivative of the corrected spectrum. If the presence of an alpha peak was ambiguous, the algorithm selects the most probable one based on the IAF detected in previous time windows. See Grosselin et al., 2018 for more details ^66^.

All participants were instructed at the protocol explanation to close their eyes during the recordings. This instruction was reminded audibly at the beginning of each calibration. They were also instructed to be relaxed and try to reduce the auditory feedback volume throughout the exercises of different sessions. Previous research showed that providing no strategies yielded to better NF effects ^57^. Here, the participants were aware that the feedback volume would decrease with relaxation, but no explicit strategies were provided to them as such to allow them to reduce the auditory feedback volume; they were told to try their own strategies, which we report in the Supplementary Material as advised in the CRED-nf checklist ^17^. A relaxing landscape background (e.g. forest sounds) was played at a constant, comfortable volume during each exercise. The audio feedback was an electronic chord sound added to this background with a volume intensity derived from EEG signals. More precisely, the individual alpha amplitude was computed in consecutive 1-second epochs as the root mean square (RMS) of EEG activity filtered in IAF±1Hz band (*NF index*); it was normalized to the calibration baseline activity to obtain a 0-1 scale, which was used to modulate the intensity of the feedback sound (*V*) in the NF group. More precisely, for each session, a baseline value was obtained from alpha activity during the corresponding 30-second calibration phase without the low quality EEG segments as assessed by a dedicated algorithm (see “EEG recording and preprocessing” section above). Coefficients were applied to this baseline value in order to define the lower (*m*) and upper (*M*) thresholds of alpha activity during the session. During the NFT, *V* was varied as a reverse linear function of the individual alpha amplitude relative to these upper and lower bounds. If the individual alpha amplitude was becoming lower than *m*, then *V* was set to 1 (maximal). If an alpha amplitude beyond *M* was reached, then *V* was set to 0 (minimal). For the EEG segments detected as noisy (LOWq quality) during the preprocessing step, *V* was set to 1. For the participants in the control group, the instruction was identical but they received sham-FB, which was the replayed feedback variations from another subject randomly chosen from the NF group at the same training level (i.e. session). For instance, a participant in the control group at the 3^rd^ session received the auditory feedback generated and received by a random subject from the NF group at the 3^rd^ session.

### Data analysis

#### NF index and learning score

For each participant and each training session, we first computed the average value of the NF index (before normalization) for every exercise. Second, in order to take into account inter-subject variability at the first session for NF index (see Fig. 3a and Supplementary Fig. S13), we built an NF learning score (Δ*D(t)*)—from the NF index variations across exercises and sessions ^67^. To do this, we computed the median value (*med*) of the NF index across the 7 exercises of the first session; then, for each session *t*, we computed *D(t)*, the number of NF index values (1 by second) above or equal to this median value *med*. This cumulative duration was divided by the total duration of the training session cleaned from LOWq segments (maximum 21 minutes) in order to express *D(t)* by minute, and transformed into percent change relative to the first session, as follows (Eq. (1)):

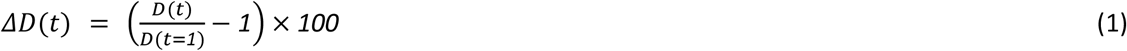

**Figure 3.**
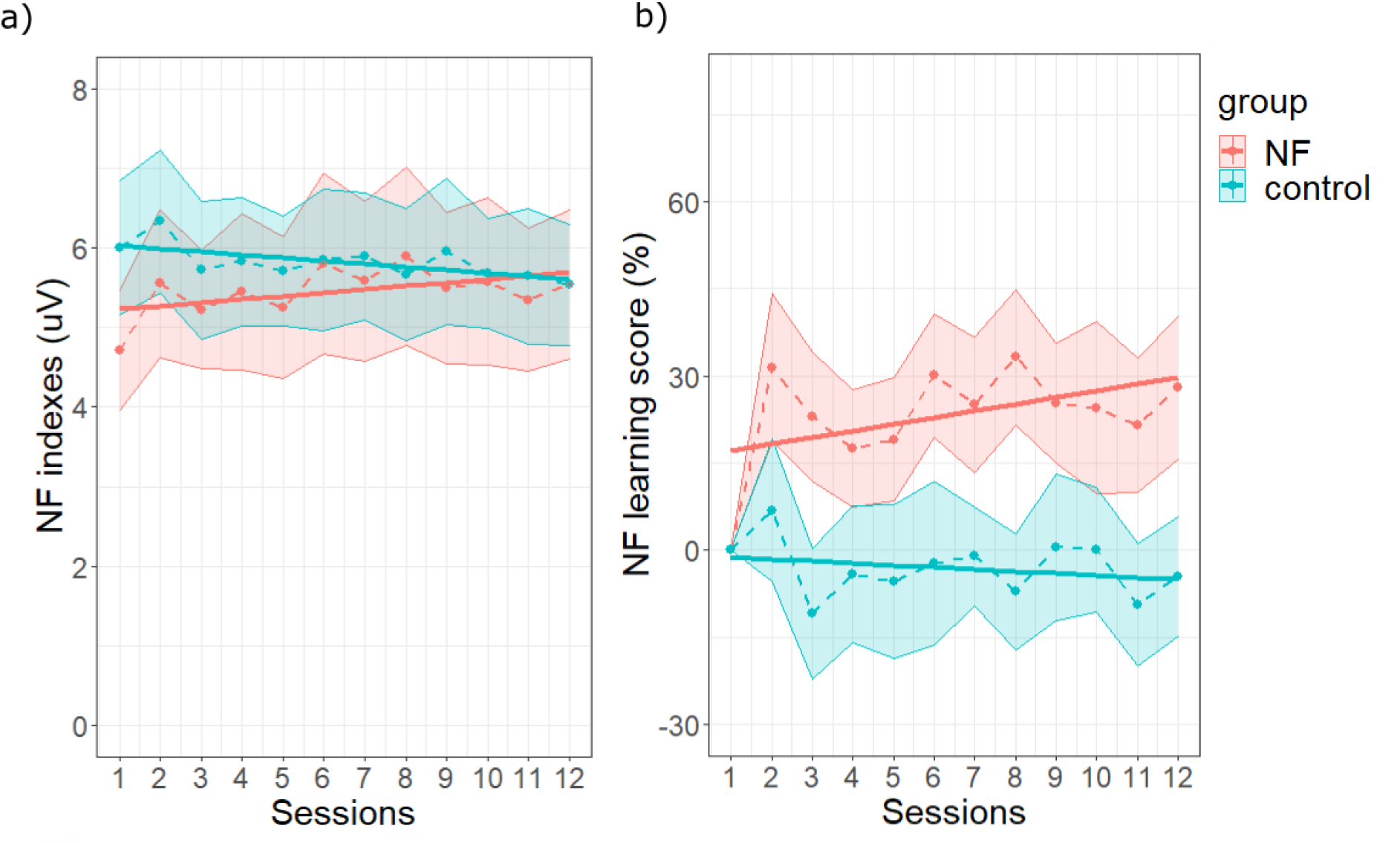
Neuromodulation induced by NFT. a) Evolution of the NF index across sessions, for the NF (in red) and the control (in blue) groups. b) Evolution of the NF learning score across sessions. In both subplots, dotted lines and points represent the values of the NF index along the 12 sessions, averaged across participants for each group; the shaded areas represent the standard errors of the means. The solid lines represent the session effect estimated for each group from the LMM.

#### Theta and low beta activities

To study the selectivity of the neuromodulation only for the targeted alpha activity, we analyzed the between-session evolutions of theta (4-7Hz) and low beta (13-18Hz) activities, as control outcomes ^67^. For each subject, on each exercise and session, theta activity was computed every second as the RMS of EEG activity filtered between 4 and 7Hz in 4-second sliding windows, on epochs with high or medium quality (see ^62^ for details about signal quality computation). We then averaged these RMS values for each session. Similar computations were performed for the EEG activity between 13 and 18Hz (low beta activity).

#### Signal quality

As encouraged in ^17^, the quality of EEG signals was analyzed to assess the poor quality EEG data prevalence between groups and across sessions. For each participant, session, and exercise, the quality of each 1-second EEG epoch recorded by each electrode was determined by a classification-based approach according to three labels: HIGHq, MEDq, and LOWq (see ^62^ for more details). A quality index *Q* was then computed for each electrode, during each exercise, as in Eq. (2):

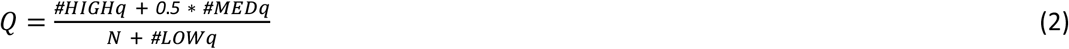

with: #HIGHq, #MEDq, #LOWq indicating the number of high, median, low quality epochs and N, the total number of quality labels during the session. Finally, the average value of *Q* was computed from the two electrodes for each exercise.

#### Self-report outcomes

The raw scores of the *STAI-Y-A* (between 20 and 80) and *relax-VAS* (between 0 and 10) were computed pre- and post-session. The subjective level of feedback control was measured within- and between-session on the *control-VAS* (between 0 and 10).

The raw scores of the *STAY-Y-B*, *PANAS* and *PSS* were obtained pre- and post-program; these latter outcomes are reported in Supplementary Material.

#### Statistical analyses

All statistical analyses were performed using R (v.4.0.2; R Core Team, 2020) and *lme4* package ^68^. We used Linear Mixed Models (LMMs) ^69,70^, because LMMs allow handling missing data, variability in effect sizes across participants, and unbalanced designs ^71^. Available data in this study are detailed in Supplementary Table S8.

For all LMM analyses, the NF group at the first session was set as the level of reference in order to specifically estimate the effects of NFT in this group. For each outcome variable studied, the choice of the random factors was done comparing the goodness of fit of the models that converged with different random factors, based on Akaike Information Criterion (AIC)^72^, Bayesian Information Criterion (BIC), log-likelihood comparison (logLik) and by running an analysis of variance (*anova*) between models. The detailed procedure for each outcome variable can be found in Supplementary Material in section 6, pages 14-20. To be concise in the main text, the random factors chosen were directly reported between parenthesis in the LMM equations below.

Similarly to ^73^, to analyze the within- and between-session NFT effects on the NF index we used fixed effects of session, exercise, group, and the 2-way interactions between session and group and between exercise and group in the following equation (Eq. (3) as coded in R, with a colon indicated an interaction between terms):

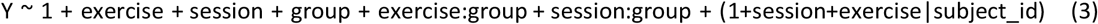

Results (see “5. NF index and feeling of control across exercises: U-curves” section in Supplementary Material pages 7-13) indicated that the effect of exercises followed a U-curve. Therefore, the exercises were coded as a quadratic term, that is, exercises 1 to 7 were coded as 9, 4, 1, 0, 1, 4, and 9. The sessions were coded as a numeric variable between 0 and 11. Eq. (3) was also used for the analysis of the *control-VAS* scores with 1+session|subject_id as random effects structure.

For the analysis of NF learning score and the signal quality index, we used the following LMM equation (Eq. (4)):

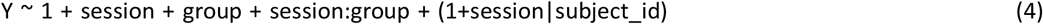

Eq. (4) was also used for the analysis of theta and low beta activities with only a random intercept by participant (1|subject_id).

For the *STAI-Y-A* outcome, we used LMM with session, phase (pre- or post-session), group, and the 2-way interactions between session and group and between phase and group as fixed effects (Eq. (5)). This model was also used for the analysis of *relax-VAS* with 1+phase|subject_id as random effects structure.

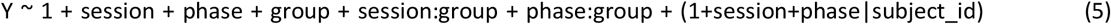

For each model, parameter *β* for the effects of interest were estimated by fitting the models on the corresponding dependent variable, using the Restricted Maximum Likelihood (REML) approach. P-values were estimated via type III Analysis of Variance on the LMM with Satterthwaite’s method, using the *anova()* function of the *lmerTest* package of R ^74^. For all variables of interest, we set *p < 0.05* as statistically significant. When there was an interaction between a factor of interest and group, LMM models were fitted in each group separately, with session and exercise or phase—as adequate—as fixed-effect factors. For these latter analyses, we used a random intercept by participant because more complex model structure generally failed to converge for at least one group ^75^. All the results of these LMM and anova analyses are reported in Supplementary Material.

Additionally, to check the variability between groups at the first session, we performed, for each outcome variable of interest, an independent t-test between groups. The results of these t-tests are presented in Supplementary Table S63.

Correlation analyses were also performed between NF index and self-report outcomes in each group as detailed in Section 13 of Supplementary Material.

## Results

### Neuromodulation induced by NF

The analysis of the NF index showed a significant interaction between session and group (F(1, 46.049) = 5.01, *p* = 0.030, supplementary Tables S25 and S26). This result suggests a significant different linear evolution of the NF index across sessions between groups. Our planned comparisons (linear mixed models run in each group) showed a significant linear increase on the NF index across sessions in the *NF group* (*β* = 0.04, CI [0.03, 0.06], F(1, 2057) = 32.43, *p < .001*, supplementary Tables S27 and S28), whereas a significant linear decrease was found in the *control group* (*β* = −0.04, CI [−0.06, −0.02], F(1, 1899) = 19.43, *p < .001*, supplementary Tables S29 and S30). These findings indicated an increase of the NF index across sessions, specific to the *NF group* (Fig. 3a). Although these results could be due to a baseline difference at the first session, an independent t-test between the mean levels of NF index of each group at session 1 did not show a significant difference (t(46) = −1.8, *p* > 0.05). See “11. Group comparison at the first session” supplementary section and supplementary Table S63 for details.

In addition, to normalize changes across sessions relative to the NF index at the first session, we built an NF learning score. This allowed us to analyze the progression of the trained activity across sessions taking into account the activity at the first session. The analysis of the NF learning score showed a marginal interaction between session and group (F(1, 46.325) = 3.27, *p* = 0.077) (see Supplementary Tables S31 and S32). Based on our *a priori* hypothesis, we looked at the session effect in each group (Supplementary Tables S33, S43, S35 and S36). The analysis of the NF learning score in each group confirmed a specific NF-based neuromodulation. Indeed, our analyses showed a significant effect of session only for the *NF group* (*β* = 1.14, CI [0.20, 2.08], F(1, 272.19) = 5.67, *p* = 0.018) (see Supplementary Tables S33 and S34), which indicates that the NF learning score increased across sessions in the *NF group* only (Fig. 3b). Additional individual linear regressions of the NF learning score (see Supplementary Fig. S8 and S9) showed that 80% (20/25) of the participants from the NF group had a positive regression slope across the 12 sessions, while the slope was positive for 48% (11/23) of the participants from the control group.

In addition, there was a significant effect of exercise on the NF index, reflecting the quadratic pattern of the NF index across exercises (F(1, 45.940) = 26.55, *p* < .001, see Supplementary Tables S2 and S3). The non-significant interaction between exercise and group indicated that this effect did not statistically differ between groups (F(1, 45.940) = 1.76e−03, *p* = 0.967) (Fig. 4).

**Figure 4.**
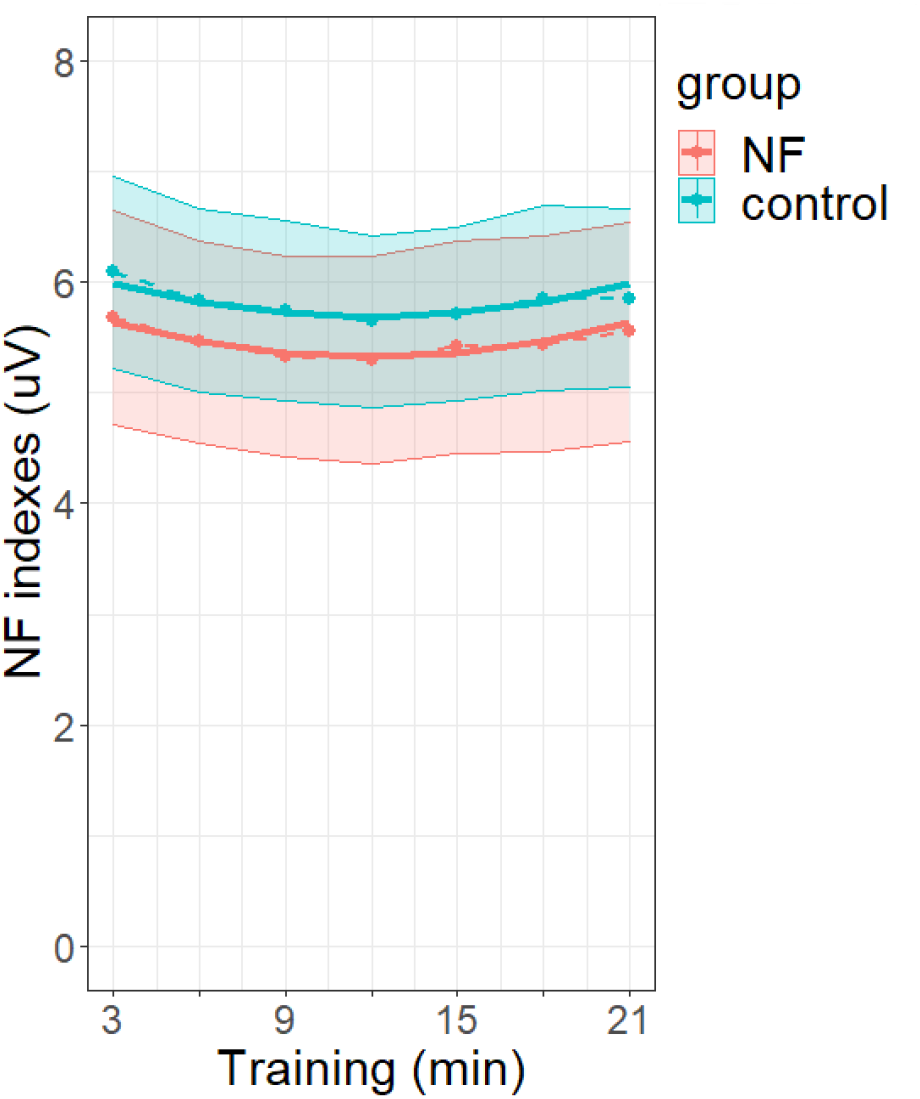
Within-sessions modulation of NF index for the NF (in red) and the control (in blue) groups. The dotted lines and points represent the values of the NF index along the training exercises, averaged across sessions and participants for each group. Training exercises are expressed in minutes (each exercise lasted 3 minutes, with 7 exercises—21 minutes training—in each session). Shaded areas represent the standard errors of the means. The solid lines represent the exercise effect estimated for each group from the LMM.

### Selectivity of the neuromodulation on alpha activity

To investigate the selectivity of the neuromodulation relative to the targeted alpha activity, we checked if the specific neuromodulation for the *NF group* occurred only for the alpha band and not for the neighboring EEG frequency bands. We analyzed EEG activity in the adjacent EEG bands, theta and low beta. For theta activity, there was a significant effect of session (F(1, 523.07) = 6.50, *p* < .011), without any statistically significant interaction between group and session (F(1, 523.07) = 2.11, *p* = 0.147). This reflected an overall increase of theta activity, with no significant difference between the NF and the control groups. For low beta activity, a marginally significant interaction between session and group was observed (F(1, 523.04) = 3.70, *p* = 0.055). However, the analyses in each group did not reveal any significant session effect either in the *NF group* (*β* = 0.01, CI [−0.01, 0.03], F(1, 272.03) = 1.40, *p* = 0.239), or in the *control group* (*β* = −0.02, CI [−0.04, 0.00], F(1, 205.01) = 2.29, *p* = 0.132). See Supplementary Tables S37, S38, S39, S40, S41, S42, S43, S44 and Supplementary Fig. S10 for details.

### Signal quality

No interaction between session and group was found on data quality *Q* (F(1, 46.130) = 0.16, *p* = 0.691), which shows no difference between groups in terms of data quality evolution across sessions. Although a marginal session effect was observed (F(1, 46.130) = 3.70, *p* = 0.061), the effect size (in terms of *β* and 95% CI) was very small (see Supplementary Tables S45 and S46 and Supplementary Fig. S11). Moreover, a chi-square analysis (see “12. Study of LOWq, MEDq and HIGHq proportions” section in Supplementary Material, page 49), showed that proportions of LOWq, MEDq and HIGHq EEG segments did not change between the first and last sessions (*X*^2^ = 0.009, df = 2, *p* = 0.9955).

### Self-report outcomes

#### Relaxation and anxiety levels

##### STAI-Y-A

The state anxiety level decreased significantly from pre- to post-session (F(1, 46.137) = 24.77, *p* < .001). This decrease did not differ statistically between groups as the interaction between phase and group was not significant (F(1, 46.137) = 2.18, *p* = 0.147) (Fig. 5a). Moreover, there was a trend to a linear decrease of *STAI-Y-A* scores across sessions (F(1, 45.787) = 3.58, *p* = 0.065), with no significant interaction between session and group (F(1, 45.787) = 1.31e−03, *p* = 0.971) (Fig. 5b; Supplementary Tables S51 and S52).

**Figure 5.**
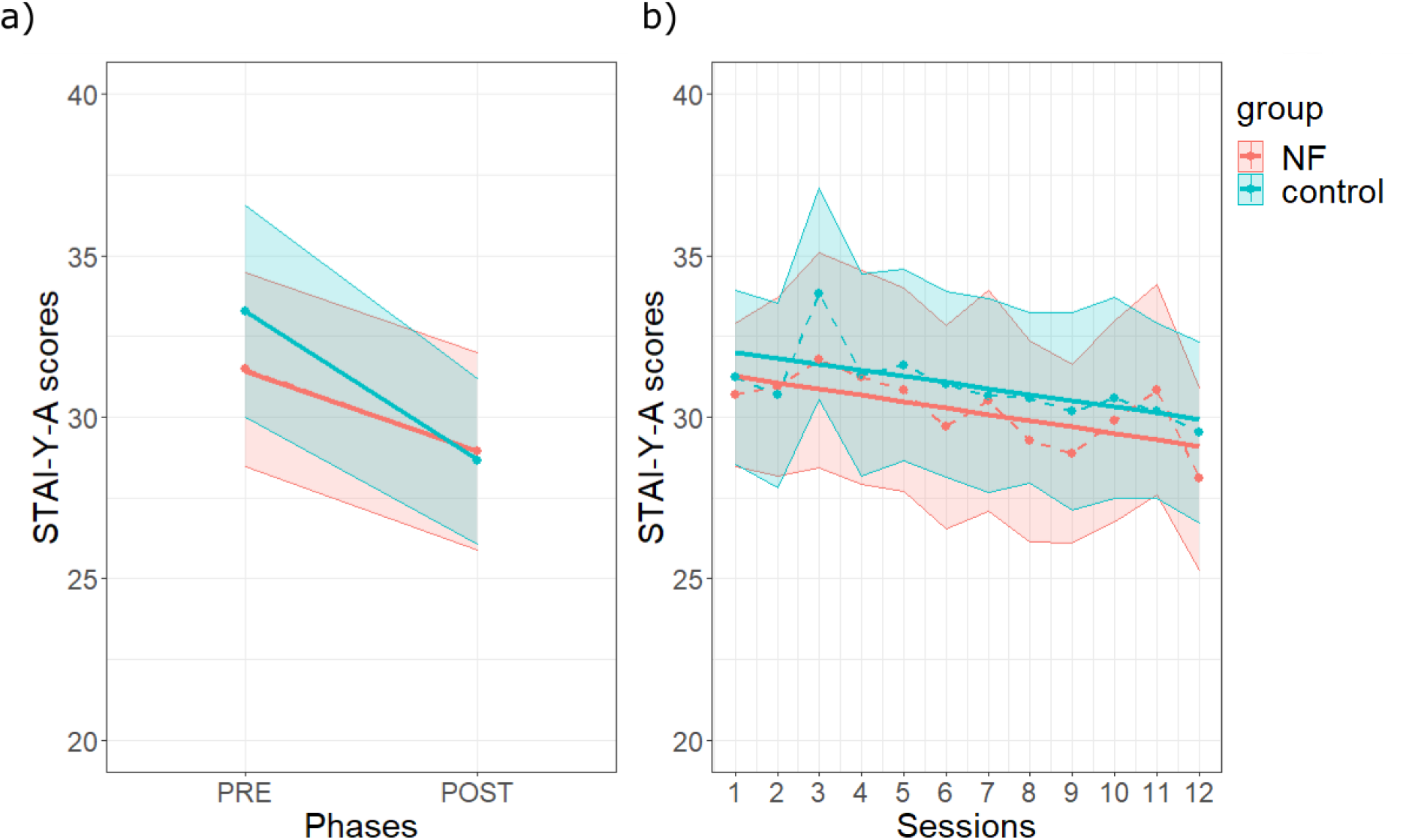
Effects of NFT on anxiety level. **a) Pre-post (phase) mean effects.** Evolution of the *STAI-Y-A* scores from pre- to post-NF training sessions, averaged across participants and sessions, for the NF (in red) and control (in blue) groups. Shaded areas represent standard errors of the means. **b) Between-session effect on *STAI-Y-A* scores in each group.** For each session, the pre- and post-session anxiety levels were averaged. Same legend as in Fig. 3a.

##### relax-VAS

Relaxation, as measured by the *relax-VAS* scores, increased from pre- to post-session (F(1, 46.01) = 34.29, *p* < .001) and this effect was not significantly different between groups (F(1, 46.01) = 0.93, *p* = 0.340) (see Fig. 6a and Supplementary Tables S53 and S54). Moreover, *relax-VAS* scores showed a significant linear increase across the sessions (F(1, 1050.22) = 18.55, *p* < .001), which did not differ significantly between groups (F(1, 1050.22) = 0.65, *p* = 0.42 (Fig. 6b; Supplementary Tables S53 and S54).

**Figure 6.**
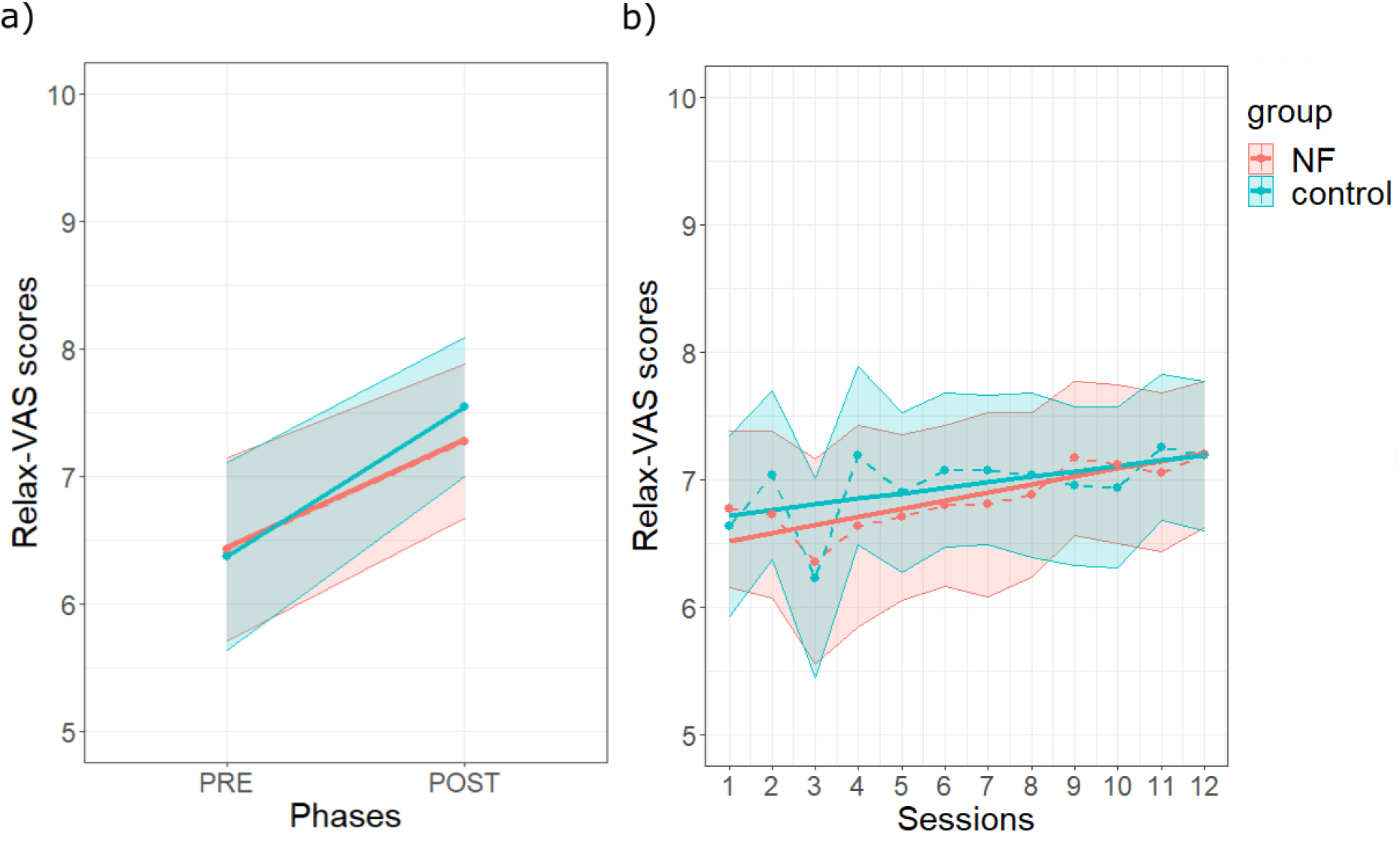
Effects of NFT on *relax-VAS* scores. **a) Pre-post (phase) mean effects. b) Between-session effect on *relax-VAS* scores in each group.** Same legend as in Fig. 5.

#### Subjective feeling of control

A significant increase of the feeling of control was observed across sessions (F(1, 46.1) = 15.40, *p* < .001). This effect was not significantly different between groups (F(1, 46.1) = 1.62, *p* = 0.209) (Fig. 7). In addition, similarly to the exercise effect on the NF index, there was a quadratic fit of control feeling over exercises (F(1, 3905.3) = 18.39, *p* < .001) which was not significantly different between groups (F(1, 3905.3) = 0.51, *p* = 0.475). See Supplementary Tables S49 and S50 and Fig. S12 for details.

**Figure 7.**
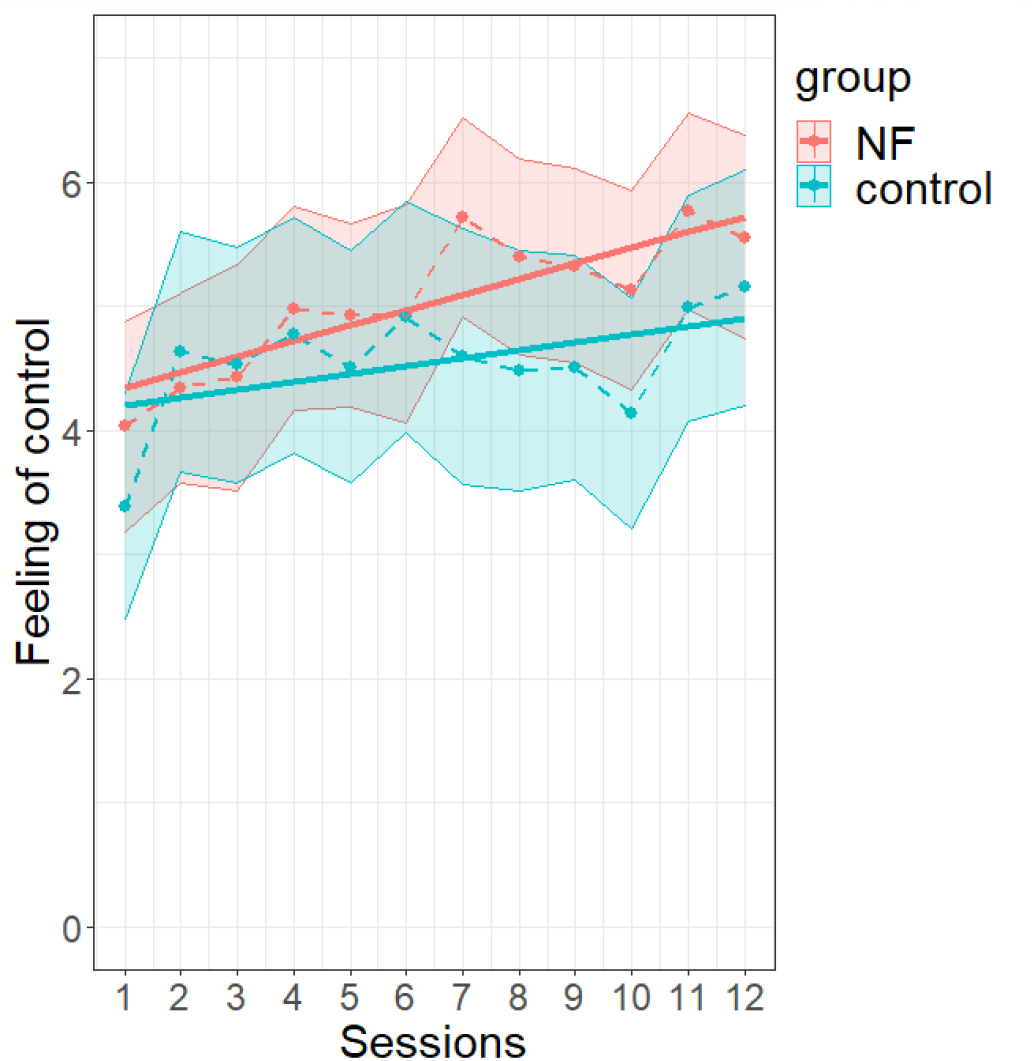
Evolution of the between-session effect of feeling of control with NF learning. See Fig. 3 for legend.

### Correlations between NF index and self-report outcomes

We examined these correlations at each session and the correlation between the slope of NF index and the slope of self-report outcomes—in terms of relaxation, anxiety, and feeling of control—across sessions (Supplementary Tables S65, S66 and S67). A few significant correlations were found at some sessions, but none of these was significant after correction for multiple comparisons. There was no significant correlation either between the slopes of NF index and self-report outcomes.

## Discussion

In this study, we proposed a double-blind sham-controlled randomized study of the neuromodulation induced by individual alpha-based NFT over 12 weekly sessions using a strictly controlled sham-FB condition as control, in healthy adult participants. NFT was performed with a wearable, dry sensors headset, which delivered intensity-modulated auditory feedback based on EEG signal amplitude in individual alpha frequency band. To avoid non-contingency between produced efforts and the resulting feedback evolution for the control group ^22^, the control condition consisted in delivering sham-FB—a feedback replayed from randomly chosen users of the NF group at the same training stage. Hence, all participants benefited from the proposed NFT experience, but only those of the NF group experienced a link between the feedback and their own alpha activity. In addition, all the participants performed the task immersed in a relaxing auditory landscape, with their eyes closed for 21 minutes, thus constituting a common reinforcer context for relaxation.

First of all, we wanted to assess the NF learning of individual alpha-band activity upregulation. NF learning refers to the capacity to self-regulate a targeted activity in the desired direction across training sessions ^34,41,42,57,76^. More specifically, we hypothesized a neuromodulation specific to the NF training, that is to say, only the NF group was expected to increase individual alpha activity across training sessions^77^. Even if the averaged values of NF index were similar at the 12th session in both groups, our analyses of the NF index and the NF learning score confirm a specific session effect only in the NF group. This finding demonstrates, across the training program, a specific neuromodulation induced by the link between individual alpha activity and FB. Indeed, the use of a randomized double-blind protocol together with the strict sham-FB control condition in a reinforcer context allowed us to control for different confounding factors which may contribute to NFT effects. In particular, it allowed controlling for context, task, reward, and performance, avoiding potential motivational biases in NF versus control conditions ^22^. One may wonder if training another frequency band could have constituted an alternative sham condition for the control group. However, as mentioned in Introduction, such control condition may induce incongruity between task instruction and the target activity in the control condition. This could render the task more difficult or less rewarding for the subject in the control condition, due to this incongruity. This is why we chose the present yoked feedback.

We observe that the NF index seemed different between NF and control groups at the first session. This could be explained by the inter-subject variability of alpha rhythm ^78^. We tested this difference as well as that of other outcome variables in the first session, and it was found to be not significant (*p*>0.05; c.f. Supplementary Table S63), showing that there was no statistically significant difference between groups at the first session for any of the studied variables. Similarly, one may note that NF index values seemed similar between groups at the 12th, final session. However, the important inter-individual variability in alpha activity makes it important to consider across sessions effects, as we did in our analyses, rather than NF index value at either first or final session.

In this study, we also examined if the neuromodulation was selective of the targeted activity ^79–81^. As proposed in ^67^, we analyzed two adjacent frequency bands (theta and low beta). We found an overall increase across sessions for the theta band, not specific to the NF group, and no significant change for the low beta band. This indicated that the neuromodulation was selective of the alpha activity in the sense that there was an absence of evidence for similar effects in the theta and beta bands.

To the best of our knowledge, this is the first evidence of selective longitudinal alpha-band neuromodulation (over 12 weeks) in a double-blind randomized study implying healthy participants trained with a wearable dry sensors NF device.

Interestingly, we could not assess an alpha neuromodulation within sessions (across exercises) as one may have expected it ^34,42,76^. In fact, we observed a U-shape pattern for the dynamics of alpha band activity during NF learning across exercises. To observe the NF learning effects, multiple sessions are required ^41,82^, in order for the participants to find their own strategies to succeed in the task ^57^. In contrast, the effects observed within-session may not only be related to relaxation training but also to other processes put at play during each session. Alpha activity is a spontaneous but complex rhythm associated with several cognitive states and processes. Its modulation has been predominantly related to vigilance, attention ^83,84^, awake but relaxed state ^50,55,56,85–89^. The alpha activity change across exercises during the sessions could reflect the different cognitive processes involved by the task: continuous monitoring of the feedback may have required heightened focused attention ^90,91^, error detection ^92,93^, and working memory processes ^94^ during the first training minutes, allowing participants to progressively adapt their cognitive strategy and mental state to the task. It is important to note that the within-session U-shape pattern of alpha activity was observed in both the NF and control groups. This supports the idea that the sham-FB condition allowed us to rigorously control for the task performed by the subjects. Altogether, the specific neuromodulation of alpha activity induced by NFT was revealed only in the longitudinal effect across the twelve sessions.

We also examined the possible effect of EEG data quality on NF learning ^17^. EEG quality did not change significantly across sessions in either group. Thus, it was not found to contribute to the neuromodulation observed in the NF group. One may wonder if we monitored the compliance of keeping eyes closed during the recordings because of the potential effects of eyes open and eyes closed on alpha activity. Even though this instruction was reminded auditorily at the beginning of each calibration, it could not be checked for the 28 participants who underwent the NFT sessions at home or at work. It has to be noted that if the participants didn’t respect this instruction during the calibration or the training sessions, this may have had an impact on data quality, hence on the feedback. For future experiments, it will be interesting to find a way to monitor this aspect of the task.

In this study, we were also interested to know if self-report benefits would be induced by the NFT and if a difference would be found between groups knowing the common reinforcer (relaxing) context of the protocol. We investigated the self-report changes in terms of relaxation and anxiety levels pre- and post-session and across the training program. We found significant benefits in terms of relaxation and anxiety from pre- to post-session, as well as a slight reduction of anxiety level and a significant increase of relaxation across sessions, but without any group difference. This can be explained by the use of sham-FB condition and the NF task proposed in our protocol, which could produce the same immersive, relaxing experience in the participants of both the control group and the NF group.

There was no significant correlation between NF index and self-report outcomes either. Thus, the self-reported benefits were not specific to the NF operant learning itself but may be explained by non-specific mechanisms of the NFT, such as the psychosocial factors (like education level, locus of control in dealing with technology, capacity to be mindful, field of work, etc.) ^30,31,33^, relaxing training context, the instructions (closed eyes during a break of 21 minutes), and repetition-related effect ^17^, which fuels the debate about the placebo effect of the neurofeedback ^30,59^. Note that education levels and the professions of the participants had almost the same repartition in both groups as well as the frequency of practice of meditation, sophrology, relaxation, arts (see Supplementary Tables S2, S3, S4, S5, S6 and S7). Furthermore, we must notice that all the subjects involved in our study were ranked as low to moderately anxious, which might have contributed to the lack of difference between groups. Indeed, Hardt and Kamiya ^95^, in their alpha-upregulation NFT study, observed reduction of anxiety level for high but not low anxious subjects. Further investigations with high anxious or clinical participants should allow to test if benefits in terms of relaxation and anxiety may be highlighted specifically for the NF group.

Overall, our findings showed that NFT induced positive self-report benefits for all participants, but the part due to neuromodulation remained unclear. Indeed, the links between self-report outcomes and neurophysiology are complex and include several factors ^17^, such as cognition, attention, motivation ^33^, training frequency ^82^, but also the choice of the neuromarker itself ^34,58^. In this study, we chose, as in most NF protocols aiming at anxiety reduction, stress-management or well-being ^14,15,55,56^, to use alpha activity as a biomarker for its known link with relaxed or meditative states ^35,43–49^. However, the alpha activity is not the unique biomarker of stress management, anxiety, relaxation and well-being. For instance, it can be a marker for attention ^90,91^ or memory ^94^. In addition, other biomarkers such as theta activity ^96–98^, beta activity ^99,100^ or the ratio theta/alpha ^43,45^ have also been associated with stress and/or anxiety reduction. Such biomarkers could be interesting targets to investigate in order to optimize our NF protocol. Further investigations should focus on the research of specific biomarkers related to psychophysiological factors, for example using neurophenomenology to study the link between neural activity modulation and participant’s inner experience ^101^.

Finally, to study the effect of sham-FB, we asked participants to assess their feeling of control during the training ^102^. We found an increase of the feeling of control across sessions in both groups, which suggests that participants of the control group were not aware of the non-contingency between their efforts and the feedback signal and had a qualitatively similar experience as those of the NF group. Although the increase in the feeling of control across sessions seemed more marked in the NF group, there was no significant difference between groups on this outcome variable. This emphasizes the closely controlled nature of our sham control condition. It suggests that our manipulation of the sham feedback remained fully implicit to the subjects. One may note that we did not check the locus of control of the participants in dealing with technology, which may have an impact on the training ^33^.

To conclude, our study demonstrated an upregulation of the individual alpha-band activity specific to the NF group with a wearable dry-sensor EEG device across multiple sessions of NF training. In contrast, self-reported effects in terms of relaxation and anxiety were observed in both the NF and the control group. Even if the relationship between the targeted EEG modulation and self-report outcomes is complex and remains to be fully elucidated, this study with a wearable dry-sensor EEG device underlined that NF can be used outside the lab to investigate and generalize NF learning mechanisms in ecological context.

## Data availability

The datasets generated during the current study are not publicly available due to the subject’s consents and restrictions of the ethics protocol to protect the privacy of subjects involved in the study.

## Acknowledgments

The authors thank myBrain Technologies for providing the dry wearable mobile EEG headsets used in this study. The authors thank the Electronic Department of myBrain Technologies, Paris, France, in particular N. Pourchier and M. Bensoussan, for their contribution to the development and improvement of the electronic part of the EEG device, as well as S. Zecri for the development of the mobile application used in this study. We thank C. Jeunet and S. Baillet for their advice on the manuscript. We thank M. Chaumon for his advice regarding a *posteriori* estimation of statistical power. Finally, the authors thank all the participants for their participation in this study.

## Author Contributions

**F.G.:** Conceptualization, Data curation, Formal analysis, Investigation, Writing - Original Draft, Writing - Review & Editing, Visualization, Project administration. **A.B.:** Conceptualization, Methodology, Formal analysis, Investigation, Writing - Original Draft, Writing - Review & Editing. **L.Y.-C.:** Formal analysis, Writing - Review & Editing. **X.W.:** Data curation, Formal analysis, Investigation, Writing - Review & Editing. **G.S.:** Investigation, Writing - Review & Editing. **L.H.:** Investigation, Resources, Writing - Review & Editing. **P.F.:** Conceptualization, Methodology, Writing - Review & Editing. **Y.A.:** Conceptualization, Methodology, Writing - Review & Editing, Supervision. **X.N.-S.:** Conceptualization, Investigation, Writing - Original Draft, Writing - Review & Editing, Project administration. **M.C.:** Methodology, Investigation, Writing - Review & Editing. **N.G.:** Conceptualization, Methodology, Writing - Original Draft, Writing - Review & Editing, Supervision, Project administration.

