## Supplementary Material for "Alpha activity neuromodulation induced by individual alpha-based neurofeedback learning in ecological context: A double-blind randomized study"

#### 1. Descriptive data

Table S1 **Descriptive data on this study**

| | **Control group** | | | **NF group** | | |
|---|---|---|---|---|---|---|
| **Nb participants** | 23 | | | 25 | | |
| **Gender** | 11 women/12 men | | | 13 women/12 men | | |
| | *Mean* | *SD* | *Min-Max* | *Mean* | *SD* | *Min-Max* |
| **Age** | 32.9 | 10.7 | 18-56 | 33.6 | 10.9 | 20-60 |
| **NF index (µV) across sessions** | 5.7 | 2.53 | 1.88-16.09 | 5.39 | 2.95 | 1-18.27 |
| ***STAI-Y-A* score in PRE session** | 33.26 | 10.27 | 20-63 | 31.48 | 9.49 | 20-71 |
| ***STAI-Y-A* score in POST session** | 28.64 | 8.12 | 20-53 | 28.97 | 9.66 | 20-71 |
| ***relax-VAS* score in PRE session** | 6.37 | 2.32 | 0.9-10 | 6.42 | 2.28 | 0.15-9.95 |
| ***relax-VAS* score in POST session** | 7.54 | 1.75 | 1.4-10 | 7.27 | 1.91 | 0.1-10 |
| ***STAI-Y-A* score before the program** | 33.48 | 8.6 | 21-51 | 32.2 | 7.65 | 20-47 |
| ***relax-VAS* score before the program** | 6.20 | 2.45 | 0.9-9.95 | 6.64 | 2.07 | 2.05-9.1 |
| **Feeling of control across the program** | 4.55 | 2.99 | 0-10 | 5.04 | 2.6 | 0-10 |
| **Pre-program *STAI-Y-B*** | 41.09 | 9.28 | 25-62 | 39.6 | 8.19 | 25-56 |
| **Post-program *STAI-Y-B*** | 39.61 | 8.44 | 27-55 | 38.96 | 7.15 | 27-56 |
| **Pre-program positive affects (PANAS)** | 35.7 | 5.94 | 21-47 | 35.76 | 5.73 | 28-44 |
| **Post-program positive affects (PANAS)** | 35.65 | 7.77 | 17-50 | 33.54 | 7.06 | 18-48 |
| **Pre-program negative affects (PANAS)** | 18.48 | 6.24 | 11-30 | 18.36 | 5.98 | 10-32 |
| **Post-program negative affects (PANAS)** | 18.17 | 5.73 | 11-30 | 18.46 | 7.68 | 11-44 |
| **Pre-program PSS** | 37.57 | 7.07 | 26-54 | 37.08 | 6.47 | 22-49 |
| **Post-program PSS** | 35.87 | 8.06 | 18-53 | 37.04 | 7.24 | 24-52 |

#### 2. Socio-demographic data

Table S2 **Education level at the first session for each group.** The numbers indicate how many participants were in each condition.

| Groups | Cannot read or write | No formal education but can read | Primary education | Secondary education | University education |
|---|---|---|---|---|---|
| **control** | 0 | 0 | 0 | 4 | 19 |
| **NF** | 0 | 0 | 0 | 3 | 22 |

Table S3 **Profession category at the first session for each group.** The profession categories were the following: A) Administration, B) Art and Culture, C) Business and Support, D) Finance, E) Management, F) Medical, G) Research and Data Analysis, H) Teacher, I) Technical and Engineering development, J) Student, K) Other. The numbers indicate how many participants were in each condition.

| Groups | A | B | C | D | E | F | G | H | I | J | K |
|---|---|---|---|---|---|---|---|---|---|---|---|
| **NF** | 2 | 4 | 2 | 0 | 5 | 0 | 4 | 1 | 4 | 3 | 0 |
| **control** | 0 | 2 | 3 | 1 | 0 | 3 | 2 | 2 | 7 | 2 | 1 |

Table S4 **Practice of sport reported at the first session by the participants of each group.** The numbers indicate how many participants were in each category of sport practice (from never practicing sport to practicing sport everyday).

| Groups | Never | Rarely | Sometimes | Often | Everyday |
|---|---|---|---|---|---|
| **NF** | 4 | 0 | 1 | 18 | 2 |
| **control** | 6 | 0 | 6 | 9 | 2 |

Table S5 **Practice of music reported at the first session by the participants of each group.** The numbers indicate how many participants were in each category of music practice (from never practicing music to practicing music everyday).

| Groups | Never | Rarely | Sometimes | Often | Everyday |
|---|---|---|---|---|---|
| **NF** | 12 | 0 | 2 | 10 | 1 |
| **control** | 14 | 0 | 3 | 5 | 0 |

Table S6 **Practice of meditation/sophrology/relaxation reported at the first session by the participants of each group.** The numbers indicate how many participants were in each category of meditation/sophrology/relaxation practice (from never practicing to practicing everyday).

| Groups | Never | Rarely | Sometimes | Often | Everyday |
|---|---|---|---|---|---|
| **NF** | 16 | 1 | 5 | 3 | 0 |
| **control** | 17 | 0 | 2 | 3 | 1 |

Table S7 **Practice of art reported at the first session by the participants of each group.** The numbers indicate how many participants were in each category of art practice (from never practicing art to practicing art everyday).

| Groups | Never | Rarely | Sometimes | Often | Everyday |
|---|---|---|---|---|---|
| **NF** | 17 | 0 | 2 | 2 | 4 |
| **control** | 16 | 0 | 3 | 1 | 2 |

### 3. Reported mental strategies

At the end of each session, participants had a debriefing questionnaire in which they were asking to report the strategies used during the entire session. We decomposed the strategies as the following:

- Projection in memories: the user thought about memories that arose from listening to the landscape sound proposed during the exercise
- Body awareness: the user was focused on part(s) of his/her body or on his/her breathing or did cardiac coherence
- Visualization/Attentional focus on sounds: the subject was focused on the landscape sound (the NF indexes or the environmental sound) proposed during exercise
- Attention defocusing: the user reported to have no specific strategy; he/she thought to nothing and cleared his/her head of any thought
- Body and mental relaxation: the subject tried to relax
- Several strategies: the user used several of the previous strategies during the NF session
- Others: counting, imagination of a story not related to the landscape sound, meditation, etc.

Here, we reported the proportion of each strategy (according to the previous categorization) in each group for the entire program (Supplementary Fig. S1). From Supplementary Fig. S1, it can be seen that for both groups, the most frequently used strategies were the focus on the landscape sound and the attention defocusing.

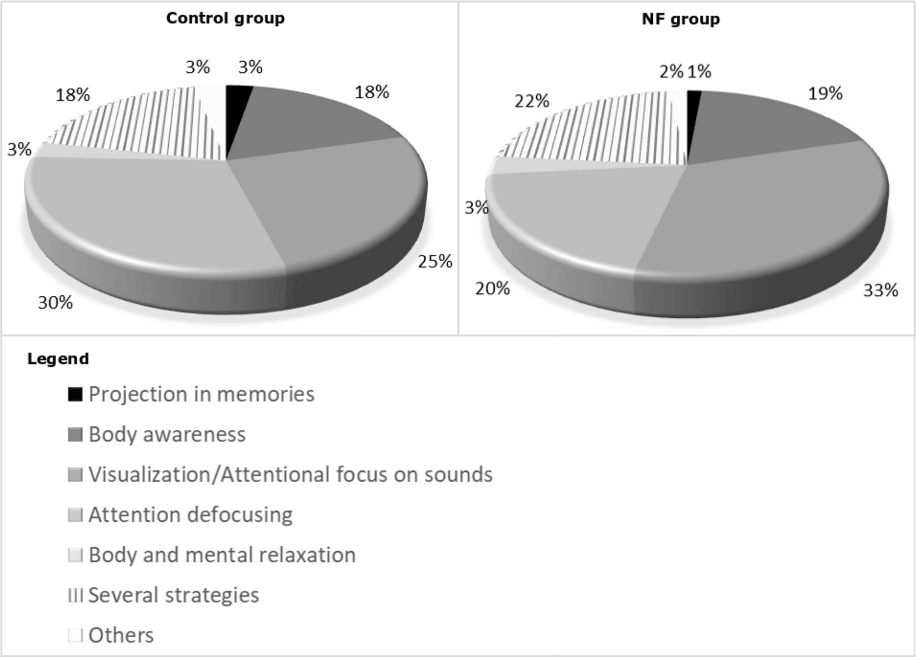


Fig. S1 **Strategies used in each group.**

### 4. Available data

Table S8 **Details of available data across subjects and the 12 NFT sessions for each group.** The number of participants in each group as well as the cumulative number of sessions are presented for the NF and control groups. The cumulative numbers of self-report questionnaires filled in across sessions and participants in each group are also presented. For each group, the percentage of achieved sessions and completed questionnaires is computed, based on the collected data and the theoretical cumulative numbers without missing data.

| | | | **Self-report questionnaires** | | | | |
|---|---|---|---|---|---|---|---|
| | **nb of users** | **nb of sessions** | ***STAI-Y-A*** | ***relax-VAS*** | ***STAI-Y-B*** | **PSS** | **PANAS** |
| **NF group** | 25 | 298 (99.3%) | 583 (97.17%) | 596 (99.33%) | 49 (98%) | 49 (98%) | 49 (98%) |
| **Control group** | 23 | 275 (99.64%) | 551 (99.8%) | 552 (100%) | 46 (100%) | 46 (100%) | 46 (100%) |

### 5. NF index and feeling of control across exercises: U-curves

#### 5.1. NF index

We first visualize, in each group, the NF index values across each session (Fig. S2 and S3). In both figures Fig. S2 and S3, we observed for most of the sessions, a quadratic progression of the NF index values across the 21-minute training drawing a U-curve.

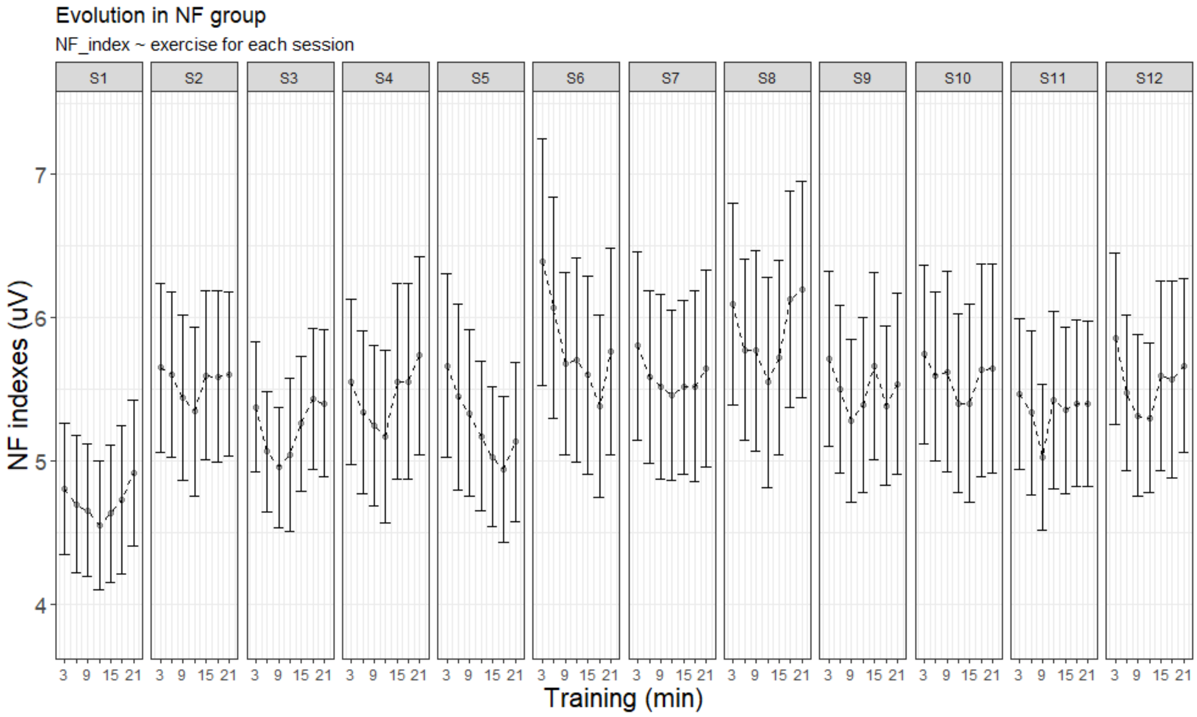


Fig. S2: **Evolution of NF index across the 21-minutes training for each session in the NF group.** The dot line represents the averaged progression of values across the 21-minute training. The error bars around the dot line is the standard error of values.

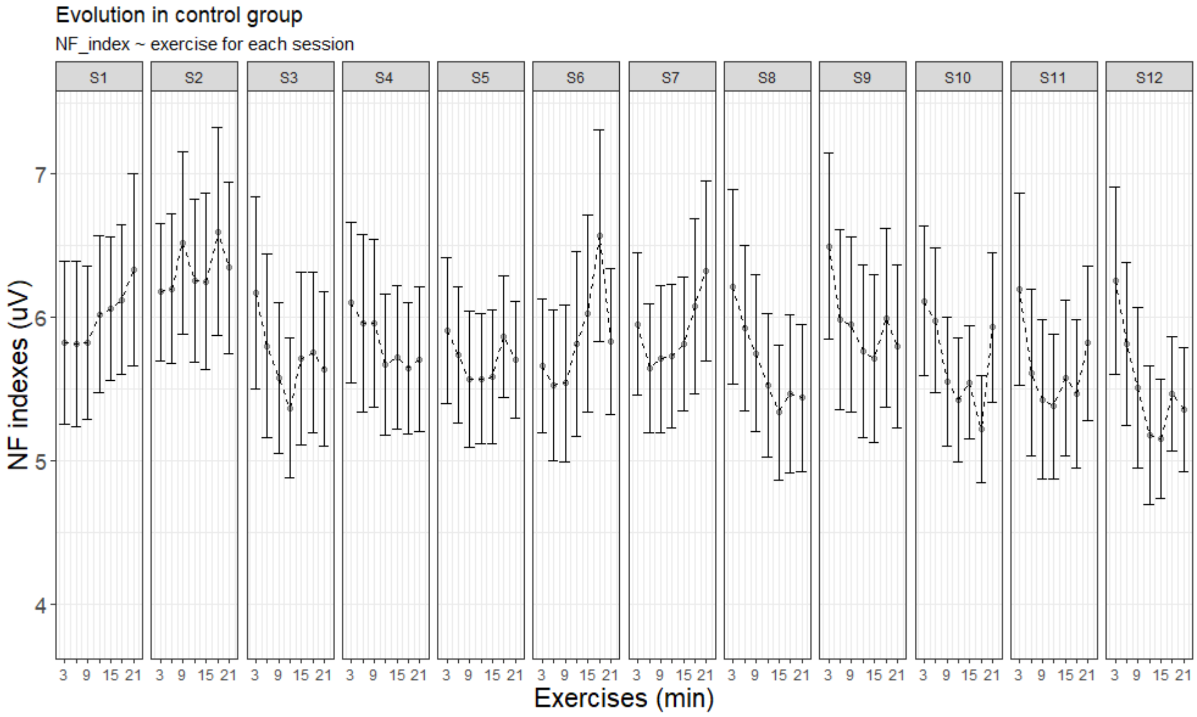


Fig. S3: **Evolution of NF index across the 21-minutes training for each session in the control group.** Same legend as in Supplementary Fig. S2.

To confirm this observation, we did two Linear Mixed Models (LMM):

- One model ($M_{lin}$) that tested the linear effect of exercises, with exercises 1 to 7 coded as 0, 1, 2, 3, 4, 5, 6
- One model ($M_{quad}$) that tested the quadratic effect of exercises, with exercises 1 to 7 coded as 9, 4, 1, 0, 1, 4, and 9.

The results obtained for each model are presented in Tables S9, S10, S11 and S12.

Table S9 **Results of $M_{lin}$ for NF index analysis.** We used an LMM including a random effect structure with random intercept by participant and fixed effects for exercise (coded as a linear term), session, group, and the two-ways interactions between exercise and group and between session and group. The NF group at the first session was set as the level of reference in order to specifically estimate the effects of NFT in this group. Thus, the parameter estimates for the effect of exercise (resp. session) corresponded to the effect of these factors in the NF group, the group[control] effect denoted the overall difference between the control and the NF group, the exercise:group [control] and session:group [control] denoted the interaction between exercise and group and between session and group respectively, estimated as the difference in parameter estimates for the exercise effect (resp. the session effect) in the control relative to the NF group. The model was fit using the Maximum Likelihood (ML) approach as we are interested in the legitimacy of a fixed effect (exercise) in the model. $\beta$ is the parameter estimate for each of the described fixed effects; 95% CI is the 95% Confidence Interval. AIC is the Akaike Information Criterion; BIC is Bayesian Information Criterion; conditional R2 is the model's total explanatory power; marginal R2 is the part of the model's explanatory power related to the fixed effects alone; Std. Dev. for standard deviation.

| | **Parameters** | $\beta$ | **95% CI** | |
|---|---|---|---|---|
| **Fixed effects** | (Intercept) | 5.25 | [ 4.27, 6.24] | |
| | **exercise** | **-9.92e-03** | **[-0.04, 0.02]** | |
| | session | 0.04 | [ 0.03, 0.06] | |
| | group [control] | 0.85 | [-0.57, 2.28] | |
| | exercise:group [control] | -0.01 | [-0.05, 0.02] | |
| | session:group [control] | -0.08 | [-0.10, -0.06] | |
| **Random effects** | **Parameters** | **Variance** | **Std. Dev.** | **Correlation** |
| | subject_id (intercept) | 6.233 | 2.497 | - |
| | residual | 1.603 | 1.266 | - |
| **Fit** | **AIC** | **BIC** | **Conditional R2** | **Marginal R2** |
| | **13559.45** | **13609.82** | 0.80 | 6.82e-03 |

Table S10 **Analysis of variance from the *$M_{lin}$* for NF index analysis**. We computed type III Analysis of Variance on the LMM of the Table S9 with Satterthwaite's method, using the *anova()* function of the *lmerTest* package of R.

| **Parameter** | **Sum Squares** | **NumDF** | **DenDF** | **Mean Square** | **F** | **p** | **Eta2 (partial)** |
|---|---|---|---|---|---|---|---|
| **exercise** | 4.82 | 1 | 3960.0 | 4.82 | 3.01 | **0.083** | 7.59e-04 |
| session | 0.22 | 1 | 3960.0 | 0.22 | 0.14 | 0.710 | 3.48e-05 |
| group [control] | 2.21 | 1 | 49.4 | 2.21 | 1.38 | 0.246 | 0.03 |
| exercise:group [control] | 0.89 | 1 | 3960.0 | 0.89 | 0.55 | 0.457 | 1.40e-04 |
| session:group [control] | 79.94 | 1 | 3960.0 | 79.94 | 49.86 | < .001 | 0.01 |

Table S11 **Results of $M_{quad}$ for NF index analysis.** The approach was identical to the one described in Table S9, except that the exercise fixed effect was coded as a quadratic term**.** See Supplementary Table S9 for table description.

| | Parameters | $\beta$ | 95% CI | |
|---|---|---|---|---|
| **Fixed effects** | (Intercept)<br>**exercise**<br>session<br>group [control]<br>exercise:group [control]<br>session:group [control] | 5.09<br>**0.03**<br>0.04<br>0.81<br>-4.63e-04<br>-0.08 | [ 4.10, 6.07]<br>**[ 0.02, 0.05]**<br>[ 0.03, 0.06]<br>[-0.61, 2.24]<br>[-0.02, 0.02]<br>[-0.10, -0.06] | |
| **Random effects** | **Parameters** | **Variance** | **Std. Dev.** | **Correlation** |
| | subject_id (intercept)<br>residual | 6.234<br>1.591 | 2.497<br>1.261 | -<br>- |
| **Fit** | **AIC** | **BIC** | **Conditional R2** | **Marginal R2** |
| | **13529.18** | **13579.55** | 0.80 | 8.35e-03 |

Table S12 **Analysis of variance from the $M_{quad}$ for NF index analysis**. We computed type III Analysis of Variance on the LMM of the Table S11 with Satterthwaite's method, using the *anova()* function of the *lmerTest* package of R.

| Parameter | Sum Squares | Num DF | DenDF | Mean Square | F | p | Eta2 (partial) |
|---|---|---|---|---|---|---|---|
| **exercise** | 53.78 | 1 | 3960.0 | 53.78 | 33.80 | **< .001** | 8.46e-03 |
| session | 0.22 | 1 | 3960.0 | 0.22 | 0.14 | 0.713 | 3.42e-05 |
| group [control] | 1.98 | 1 | 49.1 | 1.98 | 1.25 | 0.270 | 0.02 |
| exercise:group [control] | 2.58e-03 | 1 | 3960.0 | 2.58e-03 | 1.62e-03 | 0.968 | 4.09e-07 |
| session:group [control] | 79.93 | 1 | 3960.0 | 79.93 | 50.24 | < .001 | 0.01 |

These analyses showed that the linear effect of exercise was not significant, as tested in $M_{lin}$ (F(1, 3960) = 3.01, $p$ = 0.083; Table S10), whereas there was a statistically significant quadratic effect for this factor, as tested in $M_{quad}$ (F(1, 3960) = 33.8, $p$ < .001; Table S12). Moreover, lower values of AIC and BIC were observed for $M_{quad}$ than for $M_{lin}$, confirming that the $M_{quad}$ model better fitted the observed NF index values than $M_{lin}$.

#### 5.2. Feeling of control

The approach was identical to that for NF index analysis. The progression of feeling of control assessments across each session (see Fig. S4 and Fig. S5) seemed to follow the same dynamic (U-curve) as for the NF index values. To confirm this observation, we fitted the same $M_{lin}$ and $M_{quad}$ models as previously on the feeling of control values (see Tables S13, S14, S15 and S16). In Table S14, we observed that the linear effect of exercise was not significant, as tested in $M_{lin}$ ($F(1, 3955)$ = 1.71e-03, $p$ = 0.967; Table S14), whereas there was a statistically significant quadratic effect of exercise, as tested in $M_{quad}$ ($F(1, 3955) = 17.30$, $p < 0.001$; Table S16). Moreover, lower values of AIC and BIC were observed for $M_{quad}$ than for $M_{lin}$, confirming that the $M_{quad}$ model better fitted the observed NF index values than $M_{lin}$.

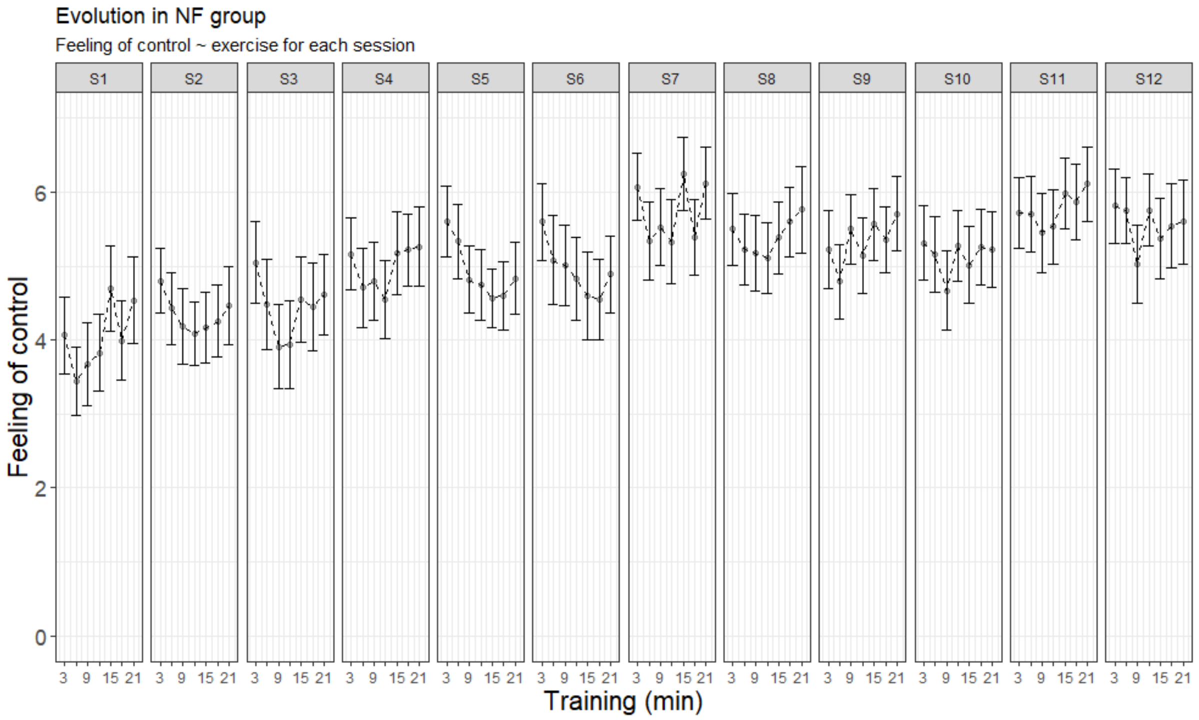


Fig. S4: **Evolution of feeling of control across the 21-minutes training for each session in the NF group.** Same legend as in Supplementary Fig. S2.

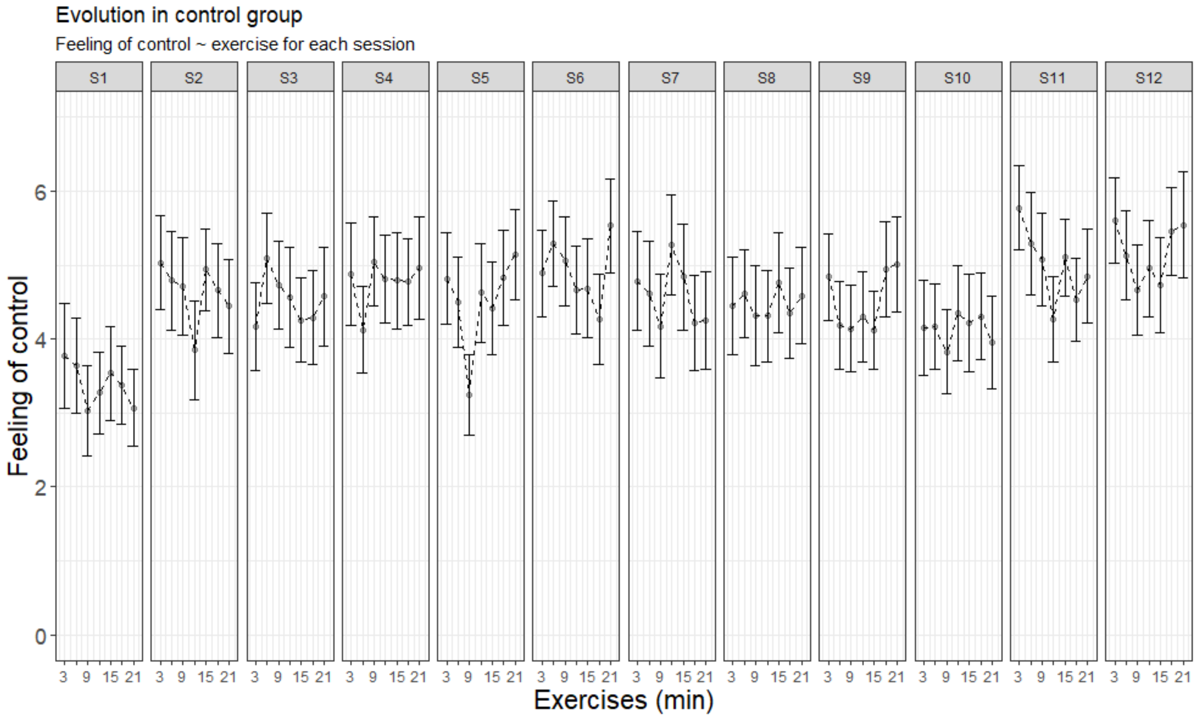


Fig. S5: **Evolution of feeling of control across the 21-minutes training for each session in the control group.** Same legend as in Supplementary Fig. S2.

Table S13 **Results of $M_{lin}$ for feeling of control analysis.** The approach was identical to the one described in Table S9**.** See Supplementary Table S9 for table description.

| | **Parameters** | $\beta$ | **95% CI** | |
|---|---|---|---|---|
| **Fixed effects** | (Intercept)<br>**exercise**<br>session<br>group [control]<br>exercise:group [control]<br>session:group [control] | 4.32<br>0.01<br>0.12<br>-0.08<br>-0.02<br>-0.06 | [ 3.57, 5.07]<br>[-0.03, 0.06]<br>[ 0.10, 0.15]<br>[-1.16, 1.00]<br>[-0.09, 0.04]<br>[-0.10, -0.02] | |
| **Random effects** | **Parameters** | **Variance** | **Std. Dev.** | **Correlation** |
| | subject_id (intercept)<br>residual | 3.349<br>4.353 | 1.830<br>2.086 | -<br>- |
| **Fit** | **AIC** | **BIC** | **Conditional R2** | **Marginal R2** |
| | **17464.54** | **17514.89** | 0.45 | 0.02 |

Table S14 **Analysis of variance from the $M_{lin}$ for feeling of control analysis**. We computed type III Analysis of Variance on the LMM of the Table S13 with Satterthwaite's method, using the *anova()* function of the *lmerTest* package of R.

| Parameter | Sum Squares | Num DF | DenDF | Mean Square | F | p | Eta2 (partial) |
|---|---|---|---|---|---|---|---|
| **exercise** | 7.44e-03 | 1 | 3955.0 | 7.44e-03 | 1.71e-03 | **0.967** | 4.32e-07 |
| session | 417.72 | 1 | 3955.2 | 417.72 | 95.96 | < .001 | 0.02 |
| group [control] | 0.09 | 1 | 55.3 | 0.09 | 0.02 | 0.883 | 3.92e-04 |
| exercise:group [control] | 2.33 | 1 | 3955.0 | 2.33 | 0.54 | 0.464 | 1.36e-04 |
| session:group [control] | 41.71 | 1 | 3955.2 | 41.71 | 9.58 | 0.002 | 2.42e-03 |

Table S15 **Results of $M_{quad}$ for feeling of control analysis.** The approach was identical to the one described in Table S9**.** See Supplementary Table S9 for table description.

| | Parameters | $\beta$ | 95% CI | |
|---|---|---|---|---|
| **Fixed effects** | (Intercept) | 4.17 | [ 3.42, 4.91] | |
| | **exercise** | **0.05** | **[ 0.02, 0.07]** | |
| | session | 0.12 | [ 0.10, 0.15] | |
| | group [control] | -0.10 | [-1.17, 0.98] | |
| | exercise:group [control] | -0.01 | [-0.05, 0.02] | |
| | session:group [control] | -0.06 | [-0.10, -0.02] | |
| **Random effects** | **Parameters** | **Variance** | **Std. Dev.** | **Correlation** |
| | subject_id (intercept) | 3.349 | 1.830 | - |
| | residual | 4.334 | 2.082 | - |
| **Fit** | **AIC** | **BIC** | **Conditional R2** | **Marginal R2** |
| | **17447.05** | **17497.41** | 0.45 | 0.02 |

Table S16 **Analysis of variance from the $M_{quad}$ for feeling of control analysis**. We computed type III Analysis of Variance on the LMM of the Table S15 with Satterthwaite's method, using the *anova()* function of the *lmerTest* package of R.

| Parameter | Sum Squares | Num DF | DenDF | Mean Square | F | p | Eta2 (partial) |
|---|---|---|---|---|---|---|---|
| **exercise** | 74.98 | 1 | 3955.0 | 74.98 | 17.30 | **< .001** | 4.36e-03 |
| session | 417.41 | 1 | 3955.2 | 417.41 | 96.31 | < .001 | 0.02 |
| group [control] | 0.14 | 1 | 53.8 | 0.14 | 0.03 | 0.858 | 6.02e-04 |
| exercise:group [control] | 2.25 | 1 | 3955.0 | 2.25 | 0.52 | 0.472 | 1.31e-04 |
| session:group [control] | 41.80 | 1 | 3955.2 | 41.80 | 9.64 | 0.002 | 2.43e-03 |

### 6. Choice of the random effects structure for the LMMs

We tried to find a compromise between parsimonious modelling of the data according to our hypotheses and the inclusion of maximal random effect structure to reduce the incidence of type I and II errors and reduce the chance of overconfident estimates (Schielzeth and Forstmeier, 2009, Barr et al., 2013; Heisig & Schäffer, 2018). Therefore, for each outcome variable, we adopted a step-wise approach to test the interest of the different random factors in the models in terms of fit to the data. As these tests concerned the random effects, we performed linear mixed models (LMM) using the Restricted Maximum Likelihood (REML) approach. Fitting random slopes in addition to a random intercept sometimes induced convergence problems because models with more complex structures need large sample sizes. When such convergence issue arises, fitting only a random intercept is better than not including random variables at all (Grueber et al. 2011). Therefore, in our approach, only models that converged were selected and they were compared to each other according to the Akaike Information Criterion (AIC) criterion (Wagenmakers & Farrell, 2004), Bayesian Information Criterion (BIC), log-likelihood comparison (logLik) and by running analysis of variance (anova) between them. For each outcome variable, we tested different random structures in this way, as detailed in each of the following subsections.

When we observed—in the final selected model—a significant or marginal interaction between fixed factors including the group term, we ran additional LMM in each group separately. This time, we only included a random intercept to account for repeated measures across sessions, because more complex model structure generally failed to converge for at least one group (Grueber et al. 2011). This procedure was applied for all the outcome variables.

#### 6.1. NF index

For NF index, we tested and compared four models with different random structures (indicated between brackets in the following formulas):

M1: Y ~ 1 + exercise + session + group + exercise:group + session:group + (1|subject_id)

M2: Y ~ 1 + exercise + session + group + exercise:group + session:group + (1+session|subject_id)

M3: Y ~ 1 + exercise + session + group + exercise:group + session:group + (1+exercise|subject_id)

M4: Y ~ 1 + exercise + session + group + exercise:group + session:group + (1+session+exercise|subject_id)

The four models converged. Thus, their goodness of fit was compared based on AIC, BIC and logLik criteria and by running the anova between models, as summarized in Table S17.

Table S17 **Comparison of models with different random structures to fit the NF index data**. npar is the number of parameters; AIC is the Akaike Information Criterion; BIC is the Bayesian Information Criterion; logLik is the log-likelihood value; P-values (Pr) were estimated via Chi-square tests (Chisq); Df are degrees of freedom of the Chi-square distribution.

| Models | npar | AIC | BIC | logLik | Chisq | Df | Pr(>Chisq) |
|---|---|---|---|---|---|---|---|
| **M1** | 8 | 13529 | 13580 | -6756.6 | | | |
| **M2** | 10 | 13214 | 13276 | -6596.8 | 319.63 | 2 | **< 0.001** |
| **M3** | 10 | 13526 | 13589 | -6753.2 | 0.00 | 0 | |
| **M4** | 13 | 13208 | 13290 | -6591.1 | 324.34 | 3 | **< 0.001** |

Thus, the M2 and M4 models statistically fitted the data better than the other models. Based on AIC and logLik criteria, the model with the maximal random effect structure was chosen (M4). Thus, the final model used for NF index analysis was the following:

**Y ~ 1 + exercise + session + group + exercise:group + session:group + (1+session+exercise|subject_id)**

The same procedure was applied for the other outcome variables as described in the next subsections.

#### 6.2. NF learning score

For NF learning score, we tested and compared two models with different random structures as indicated between brackets:

M1: Y ~ 1 + session + group + session:group + (1|subject_id)

M2: Y ~ 1 + session + group + session:group + (1+session|subject_id)

The two models converged and their goodness of fit was compared based on AIC, BIC and logLik criteria and by running the anova between models. Table S18 shows the results of this comparison.

Table S18 **Comparison of models with different random structures to fit the NF learning score data**. See Supplementary Table S17 for table description.

| Models | npar | AIC | BIC | logLik | Chisq | Df | Pr(>Chisq) |
|---|---|---|---|---|---|---|---|
| **M1** | 6 | 5561.4 | 5587.5 | -2774.7 | | | |
| **M2** | 8 | 5553.1 | 5587.9 | -2768.6 | 12.293 | 2 | **0.002141** |

The M2 model statistically fitted the data better than the M1 model. Thus, the final model used for the NF learning score analysis was the following:

**Y ~ 1 + session + group + session:group + (1+session|subject_id)**

##### 6.3. Theta activity

For the theta activity, we tested and compared two models with different random structures as indicated between brackets:

M1: Y ~ 1 + session + group + session:group + (1|subject_id)

M2: Y ~ 1 + session + group + session:group + (1+session|subject_id)

The M2 model did not converge. Therefore, we kept the M1 model. Thus, the final model used for the theta activity analysis was the following:

**Y ~ 1 + session + group + session:group + (1|subject_id)**

##### 6.4. Low beta activity

For low beta activity, we tested and compared two models with different random structures as indicated between brackets:

M1: Y ~ 1 + session + group + session:group + (1|subject_id)

M2: Y ~ 1 + session + group + session:group + (1+session|subject_id)

The two models converged and their goodness of fit was compared based on AIC, BIC and logLik criteria and by running the anova between models. Table S19 shows the results of this comparison.

Table S19 **Comparison of models with different random structures to fit the low beta data**. See Supplementary Table S17 for table description.

| Models | npar | AIC | BIC | logLik | Chisq | Df | Pr(>Chisq) |
|---|---|---|---|---|---|---|---|
| **M1** | 6 | 1167.4 | 1193.5 | -577.70 | | | |
| **M2** | 8 | 1171.3 | 1206.2 | -577.68 | 0.0444 | 2 | 0.978 |

The M2 model did not fit the data better than the more parsimonious M1 model. Thus, the final model used for the low beta activity analysis was the following (M1):

**Y ~ 1 + session + group + session:group + (1|subject_id)**

##### 6.5. Quality index

For the quality index, we tested and compared two models with different random structures as indicated between brackets:

M1: Y ~ 1 + session + group + session:group + (1|subject_id)

M2: Y ~ 1 + session + group + session:group + (1+session|subject_id)

The two models converged and their goodness of fit was compared based on AIC, BIC and logLik criteria and by running the anova between models. Table S20 shows the results of this comparison.

Table S20 **Comparison of models with different random structures to fit the quality index data**. See Supplementary Table S17 for table description.

| Models | npar | AIC | BIC | logLik | Chisq | Df | Pr(>Chisq) |
|---|---|---|---|---|---|---|---|
| **M1** | 6 | -5551.5 | -5513.7 | 2781.8 | | | |
| **M2** | 8 | -5697.9 | -5647.5 | 2856.9 | 150.38 | 2 | **< 0.001** |

The M2 model statistically fitted the data better than the M1 model. Thus, the final model used for the quality index analysis was the following:

**Y ~ 1 + session + group + session:group + (1+session|subject_id)**

##### 6.6. Timeline

To test for the effect of the time of day (timeline) at which the participants had their sessions, and to test if any difference existed between NF and control groups, we analysed the hour of the timestamp of each recording just before the beginning of the NF session.

We tested and compared two models with different random structures as indicated between brackets:

M1: Y ~ 1 + session + group + session:group + (1|subject_id)

M2: Y ~ 1 + session + group + session:group + (1+session|subject_id)

The two models converged and their goodness of fit was compared based on AIC, BIC and logLik criteria and by running the anova between models. Table S21 shows the results of this comparison.

Table S21 **Comparison of models with different random structures to fit the timeline data**. See Supplementary Table S17 for table description.

| Models | npar | AIC | BIC | logLik | Chisq | Df | Pr(>Chisq) |
|---|---|---|---|---|---|---|---|
| **M1** | 6 | 2681.3 | 2707.4 | -1334.7 | | | |
| **M2** | 8 | 2683.7 | 2718.6 | -1333.9 | 1.6075 | 2 | 0.4476 |

The M2 model did not statistically fit the data better than the more parsimonious M1 model. Thus, the final model retained for the timeline analysis was the following:

**Y ~ 1 + session + group + session:group + (1|subject_id)**

#### 6.7. Feeling of control

For the feeling of control, we tested and compared four models with different random structures as indicated between brackets:

M1: Y ~ 1 + exercise + session + group + exercise:group + session:group + (1|subject_id)

M2: Y ~ 1 + exercise + session + group + exercise:group + session:group + (1+session|subject_id)

M3: Y ~ 1 + exercise + session + group + exercise:group + session:group + (1+exercise|subject_id)

M4: Y ~ 1 + exercise + session + group + exercise:group + session:group + (1+session+exercise|subject_id)

M3 and M4 did not converge. Thus, we only compared M1 and M2 based on AIC, BIC and logLik criteria and by running an anova between these models. Table S22 shows the results of this comparison.

Table S22 **Comparison of models with different random structures to fit the feeling of control data**. See Supplementary Table S17 for table description.

| Models | npar | AIC | BIC | logLik | Chisq | Df | Pr(>Chisq) |
|---|---|---|---|---|---|---|---|
| **M1** | 8 | 17447 | 17497 | -8715.5 | | | |
| **M2** | 10 | 17285 | 17348 | -8632.3 | 166.51 | 2 | **< 0.001** |

The M2 model statistically fitted the data better than the M1 model. Thus, the final model used for the feeling of control analysis was the following:

**Y ~ 1 + exercise + session + group + exercise:group + session:group + (1+session|subject_id)**

#### 6.8. STAI-Y-A

For STAI-Y-A, we tested and compared four models with different random structures as indicated between brackets:

M1: Y ~ 1 + session + phase + group + session:group + phase:group + (1|subject_id)

M2: Y ~ 1 + session + phase + group + session:group + phase:group + (1+session|subject_id)

M3: Y ~ 1 + session + phase + group + session:group + phase:group + (1+phase|subject_id)

M4: Y ~ 1 + session + phase + group + session:group + phase:group + (1+session+phase|subject_id)

The four models converged and were compared based on AIC, BIC and logLik criteria and by running anovas between the models. Table S23 shows the results of these comparisons.

Table S23 **Comparison of models with different random structures to fit the STAI-Y-A data**. See Supplementary Table S17 for table description.

| Models | npar | AIC | BIC | logLik | Chisq | Df | Pr(>Chisq) |
|---|---|---|---|---|---|---|---|
| **M1** | 8 | 7495.8 | 7536.1 | -3739.9 | | | |
| **M2** | 10 | 7427.9 | 7478.3 | -3704.0 | 0.000 | 0 | |
| **M3** | 10 | 7424.1 | 7474.4 | -3702.1 | 75.731 | 2 | **< 0.001** |
| **M4** | 13 | 7329.9 | 7395.3 | -3652.0 | 104.027 | 3 | **< 0.001** |

The M3 and M4 models statistically fitted the data better than the M1 and M2 models. We kept the maximal random effect structure (M4) that fitted the data best. Thus, the final model used for STAI-Y-A analysis was the following:

**Y ~ 1 + session + phase + group + session:group + phase:group + (1+session+phase|subject_id)**

#### 6.9. relax-VAS

For relax-VAS, we tested and compared four models with different random structures as indicated between brackets:

M1: Y ~ 1 + session + phase + group + session:group + phase:group + (1|subject_id)

M2: Y ~ 1 + session + phase + group + session:group + phase:group + (1+session|subject_id)

M3: Y ~ 1 + session + phase + group + session:group + phase:group + (1+phase|subject_id)

M4: Y ~ 1 + session + phase + group + session:group + phase:group + (1+session+phase|subject_id)

M4 did not converge. Thus, only M1, M2 and M3 were compared based on AIC, BIC and logLik criteria and by running anova between these models. Table S24 shows the results of these comparisons.

Table S24 **Comparison of models with different random structures to fit the relax-VAS data**. See Supplementary Table S17 for table description.

| Models | npar | AIC | BIC | logLik | Chisq | Df | Pr(>Chisq) |
|---|---|---|---|---|---|---|---|
| **M1** | 8 | 4385.7 | 4426.0 | -2184.8 | | | |
| **M2** | 10 | 4373.5 | 4423.9 | -2176.7 | 0.000 | 0 | |
| **M3** | 10 | 4314.9 | 4365.3 | -2147.4 | 74.782 | 2 | **< 0.001** |

The M3 model statistically fitted the data better than M1 and M2 models. Thus, the final model used for relax-VAS analysis was the following:

**Y ~ 1 + session + phase + group + session:group + phase:group + (1+phase|subject_id)**

#### 6.10. STAI-Y-B, PANAS and PSS

For STAI-Y-B, PANAS and PSS scores, we tested and compared two models with different random structures as indicated between brackets:

M1: Y ~ 1 + phase + group + phase:group + (1|subject_id)

M2: Y ~ 1 + phase + group + phase:group + (1+phase|subject_id)

However, M2 model encountered a problem of calculation because of the number of observations (=95) that was lower than the number of random effects (=96) for the random term (1+phase|subject_id). Thus, the random-effects parameters and the residual variance were unidentifiable. For all these variables, we kept the M1 model.

### 7. Neurophysiological modulation analysis

#### 7.1. NF index

Based on the choice of the random structure, detailed in the "Choice of the random effects structure for the LMMs" section above, the following model was chosen:

Y ~ 1 + exercise + session + group + exercise:group + session:group + (1+session+exercise|subject_id)

The following table S25 presents the results of this model.

Table S25 **Results of the Linear Mixed Model (LMM) for NF index analysis.** We used the LMM described in Eq. (3) of the main text. See Supplementary Table S9 for table description.

| | | | | |
|---|---|---|---|---|
| **Fixed effects** | **Parameters** | $\beta$ | **95% CI** | |
| | (Intercept)<br>exercise<br>session<br>group [control]<br>exercise:group [control]<br>session:group [control] | 5.09<br>0.03<br>0.04<br>0.81<br>-5.47e-04<br>-0.08 | [ 4.12, 6.06]<br>[ 0.02, 0.05]<br>[-0.01, 0.09]<br>[-0.59, 2.21]<br>[-0.03, 0.03]<br>[-0.15, -0.01] | |
| **Random effects** | **Parameters** | **Variance** | **Std. Dev.** | **Correlation** |
| | subject_id (intercept)<br>session<br>exercise<br><br>residual | 6.0178081<br>0.0142493<br>0.0006175<br><br>1.4203476 | 2.45312<br>0.11937<br>0.02485<br><br>1.19178 | -<br>-0.07 (intercept)<br>[0.68 (intercept),<br>-0.33 (session)]<br>- |
| **Fit** | **AIC** | **BIC** | **Conditional R2** | **Marginal R2** |
| | 13233.64 | 13315.49 | 0.83 | 8.04e-03 |

Table S26 **Analysis of variance from the LMM of NF index**. We computed type III Analysis of Variance on the LMM of the Table S25 with Satterthwaite's method, using the *anova()* function of the *lmerTest* package of R.

| Parameter | Sum Squares | NumDF | DenDF | Mean Square | F | p | Eta2 (partial) |
|---|---|---|---|---|---|---|---|
| exercise | 37.71 | 1 | 45.940 | 37.71 | 26.55 | **< .001** | 0.37 |
| session | 9.10e-03 | 1 | 46.049 | 9.10e-03 | 6.41e-03 | 0.937 | 1.39e-04 |
| group | 1.82 | 1 | 46.004 | 1.82 | 1.28 | 0.264 | 0.03 |
| exercise:group | 2.50e-03 | 1 | 45.940 | 2.50e-03 | 1.76e-03 | 0.967 | 3.84e-05 |
| session:group | 7.12 | 1 | 46.049 | 7.12 | 5.01 | **0.030** | 0.10 |

Table S26 shows a significant interaction between group and session. Then, to examine the effect of the session within each group, we ran additional LMM in each group separately.

Table S27 **Results of the Linear Mixed Model (LMM) for NF index analysis of the NF group.** We used an LMM including a random effect structure with random intercept by participant and fixed effects for exercise and session. The model was fit using the Restricted Maximum Likelihood (REML) approach. See Supplementary Table S9 for table description.

| | | | | |
|---|---|---|---|---|
| **Fixed effects** | **Parameters** | $\beta$ | **95% CI** | |
| | (Intercept)<br>exercise<br>session | 5.09<br>0.03<br>0.04 | [3.99, 6.19]<br>[0.02, 0.05]<br>[0.03, 0.06] | |
| **Random effects** | **Parameters** | **Variance** | **Std. Dev.** | **Correlation** |
| | subject_id (intercept)<br>residual | 7.817<br>1.412 | 2.796<br>1.188 | -<br>- |
| **Fit** | **AIC** | **BIC** | **Conditional R2** | **Marginal R2** |
| | 6808.70 | 6836.91 | 0.85 | 3.85e-03 |

Table S28 **Analysis of variance from the LMM of NF index of the NF group**. We computed type III Analysis of Variance on the LMM of the Table S27 with Satterthwaite's method, using the *anova()* function of the *lmerTest* package of R.

| Parameter | Sum Squares | NumDF | DenDF | Mean Square | F | p | Eta2 (partial) |
|---|---|---|---|---|---|---|---|
| exercise<br>session | 28.40<br>45.79 | 1<br>1 | 2057<br>2057 | 28.40<br>45.79 | 20.12<br>32.43 | **< .001**<br>**< .001** | 9.68e-03<br>0.02 |

Table S29 **Results of the Linear Mixed Model (LMM) for NF index analysis of the control group.** We used the same LMM as for Table S27 but the data were those of the control group. The model was fit using the Restricted Maximum Likelihood (REML) approach. See Supplementary Table S9 for table description.

| | | | | |
|---|---|---|---|---|
| **Fixed effects** | **Parameters** | $\beta$ | **95% CI** | |
| | (Intercept)<br>exercise<br>session | 5.90<br>0.03<br>-0.04 | [ 4.97, 6.83]<br>[ 0.02, 0.05]<br>[-0.06, -0.02] | |
| **Random effects** | **Parameters** | **Variance** | **Std. Dev.** | **Correlation** |
| | subject_id (intercept)<br>residual | 5.075<br>1.788 | 2.253<br>1.337 | -<br>- |
| **Fit** | **AIC** | **BIC** | **Conditional R2** | **Marginal R2** |
| | 6725.97 | 6753.78 | 0.74 | 4.54e-03 |

Table S30 **Analysis of variance from the LMM of NF index of the control group**. We computed type III Analysis of Variance on the LMM of the Table S29 with Satterthwaite's method, using the *anova()* function of the *lmerTest* package of R.

| **Parameter** | **Sum Squares** | **NumDF** | **DenDF** | **Mean Square** | **F** | **p** | **Eta2 (partial)** |
|---|---|---|---|---|---|---|---|
| exercise | 25.49 | 1 | 1899 | 25.49 | 14.26 | **< .001** | 7.45e-03 |
| session | 34.74 | 1 | 1899 | 34.74 | 19.43 | **< .001** | 0.01 |

Additional individual linear regressions of the NF index (see Supplementary Fig. S6 and Fig. S7) showed that 68% (17/25) of the participants from the NF group had a positive regression slope across the 12 sessions, while the slope was positive for 34.8% (8/23) of the participants from the control group.

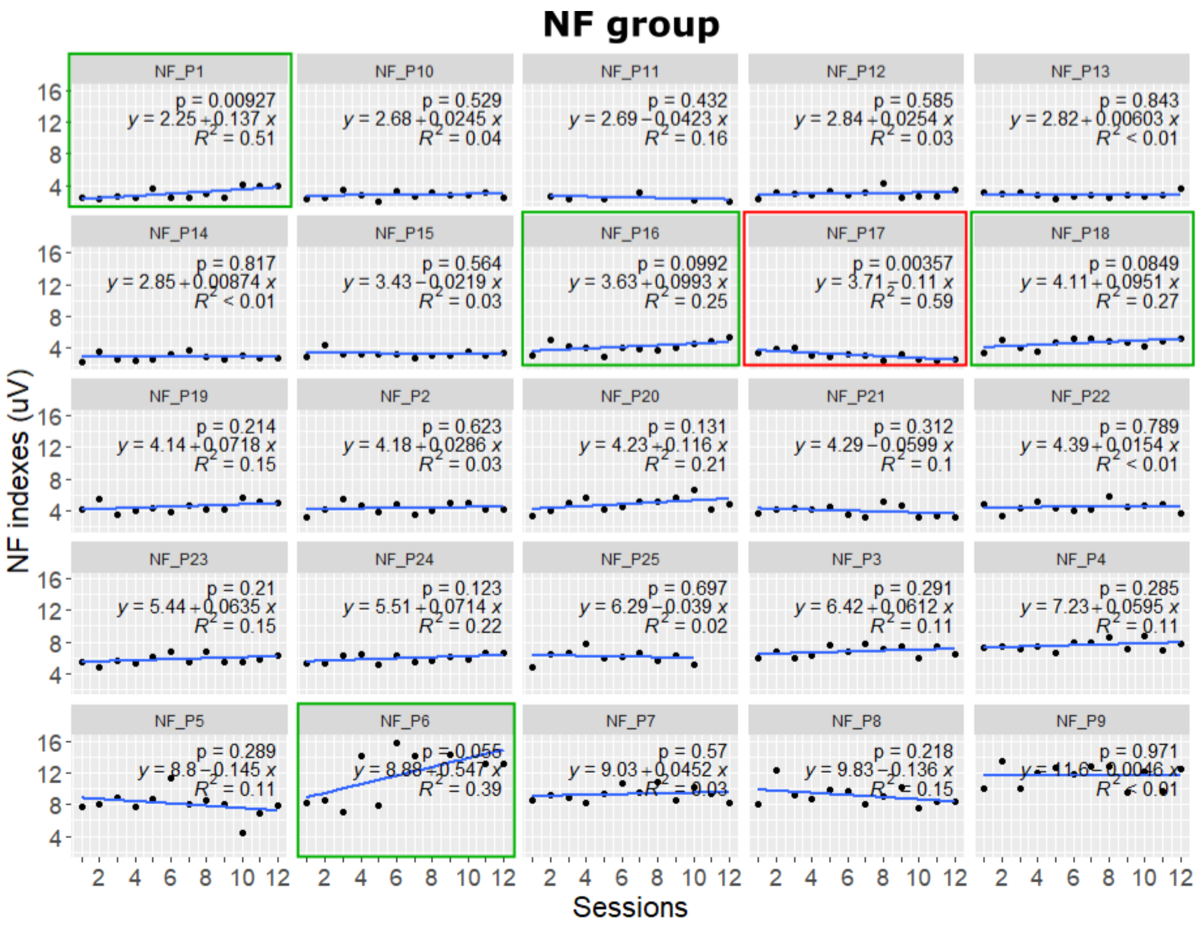


Fig. S6 **Evolution of NF index across the 12 training sessions for each participant of the NF group.** Plots of NF index values (black dots) across sessions are represented for each participant (sorted in ascending order according to the NF index value at the first session). The blue line represents the linear regression of individual NF index values across sessions of individual NF index values across sessions. The grey shaded area around the blue line is the standard error of the regression. When there is a significant or marginal increase (or decrease), the plot is framed in green (respectively, in red).

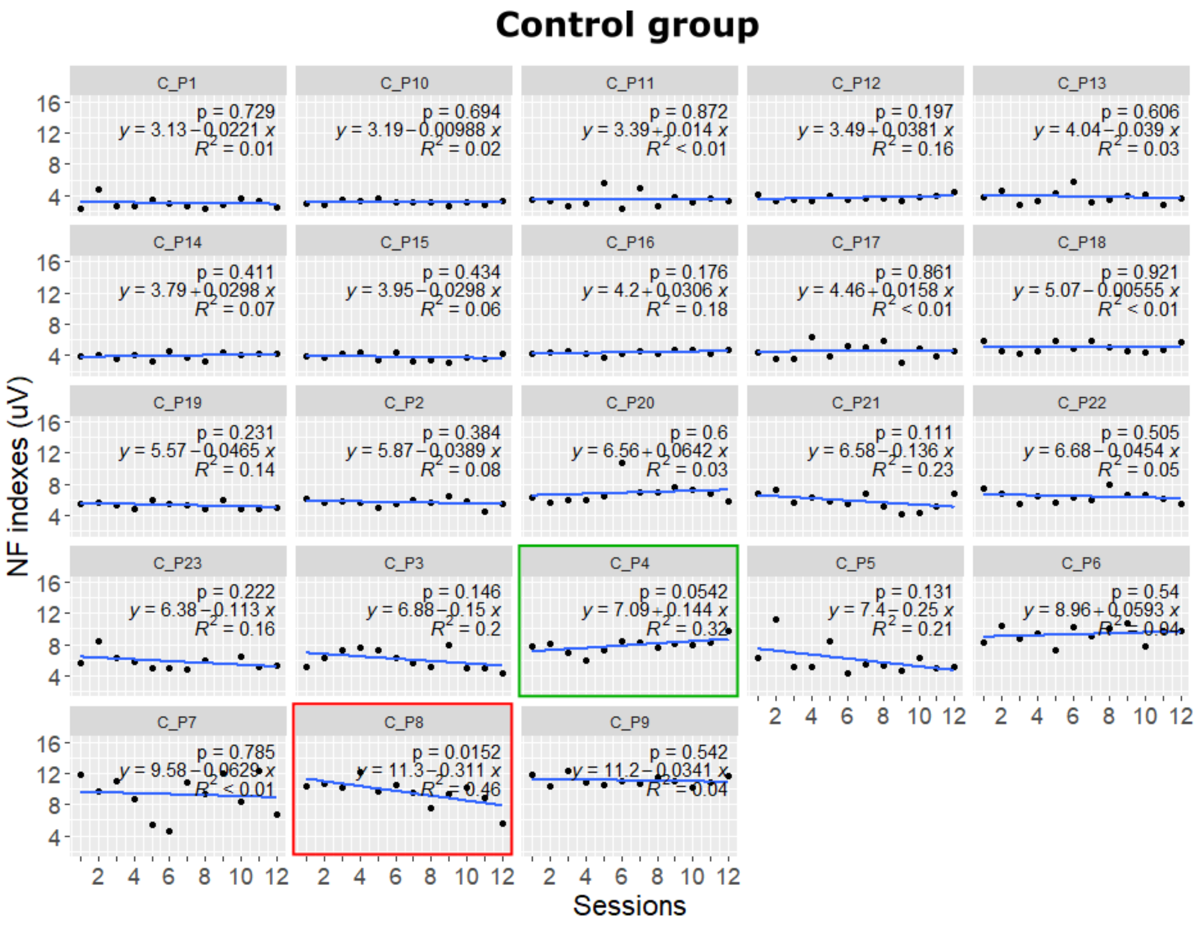


Fig. S7 **Evolution of NF index across the 12 training sessions for each participant of the control group.** Same legend as in Supplementary Fig. S6.

##### 7.2. NF learning score

Table S31 **Results of the LMM for NF learning score analysis.** We used the LMM described in Eq. (4) of the main text. The approach was identical to that for NF index analysis (described in Table S25), except that the fixed effects only included session, group and interaction between session and group**.** See Supplementary Table S9 for table description.

| | Parameters | $\beta$ | 95% CI | |
|---|---|---|---|---|
| **Fixed effects** | (Intercept)<br>session<br>group [control]<br>session:group [control] | 17.10<br>1.15<br>-18.29<br>-1.50 | [ 8.61, 25.59]<br>[ 0.02, 2.28]<br>[-30.55, -6.04]<br>[ -3.13, 0.13] | |
| **Random effects** | **Parameters** | **Variance** | **Std. Dev.** | **Correlation** |
| | subject_id (intercept)<br>session<br>residual | 242.510<br>2.882<br>763.885 | 15.573<br>1.698<br>27.638 | -<br>0.39<br>- |
| **Fit** | **AIC** | **BIC** | **Conditional R2** | **Marginal R2** |
| | 5542.25 | 5577.06 | 0.46 | 0.13 |

Table S32 **Analysis of variance from the LMM of NF learning score**. We computed type III Analysis of Variance on the LMM of the Table S31 with Satterthwaite's method, using the *anova()* function of the *lmerTest* package of R.

| Parameter | Sum Squares | NumDF | DenDF | Mean Square | F | p | Eta2 (partial) |
|---|---|---|---|---|---|---|---|
| session | 710.16 | 1 | 46.325 | 710.16 | 0.93 | 0.340 | 0.02 |
| group | 6538.35 | 1 | 45.866 | 6538.35 | 8.56 | **0.005** | 0.16 |
| session:group | 2494.13 | 1 | 46.325 | 2494.13 | 3.27 | **0.077** | 0.07 |

Table S32 shows a marginal interaction between group and session. Considering our a priori hypothesis, we performed additional LMM in each group separately.

Table S33 **Results of the Linear Mixed Model (LMM) for NF learning score analysis of the NF group.** We used an LMM including a random effect structure with random intercept by participant and fixed effects for session. The model was fit using the Restricted Maximum Likelihood (REML) approach. See Supplementary Table S9 for table description.

| | | | | |
|---|---|---|---|---|
| **Fixed effects** | **Parameters** | $\beta$ | **95% CI** | |
| | (Intercept)<br>session | 17.14<br>1.14 | [6.68, 27.60]<br>[0.20, 2.08] | |
| **Random effects** | **Parameters** | **Variance** | **Std. Dev.** | **Correlation** |
| | subject_id (intercept)<br>residual | 474.0<br>804.6 | 21.77<br>28.37 | -<br>- |
| **Fit** | **AIC** | **BIC** | **Conditional R2** | **Marginal R2** |
| | 2892.19 | 2906.98 | 0.38 | 0.01 |

Table S34 **Analysis of variance from the LMM of NF learning score of the NF group**. We computed type III Analysis of Variance on the LMM of the Table S33 with Satterthwaite's method, using the *anova()* function of the *lmerTest* package of R.

| **Parameter** | **Sum Squares** | **NumDF** | **DenDF** | **Mean Square** | **F** | **p** | **Eta2 (partial)** |
|---|---|---|---|---|---|---|---|
| session | 4566.24 | 1 | 272.19 | 4566.24 | 5.67 | **0.018** | 0.02 |

Table S35 **Results of the Linear Mixed Model (LMM) for NF learning score analysis of the control group.** We used the same LMM as for Table S33 but the data were those of the control group. The model was fit using the Restricted Maximum Likelihood (REML) approach. See Supplementary Table S9 for table description.

| | | | | |
|---|---|---|---|---|
| **Fixed effects** | **Parameters** | $\beta$ | **95% CI** | |
| | (Intercept)<br>session | -1.19<br>-0.35 | [-11.52, 9.14]<br>[ -1.32, 0.61] | |
| **Random effects** | **Parameters** | **Variance** | **Std. Dev.** | **Correlation** |
| | subject_id (intercept)<br>residual | 404.1<br>794.7 | 20.10<br>28.19 | -<br>- |
| **Fit** | **AIC** | **BIC** | **Conditional R2** | **Marginal R2** |
| | 2662.60 | 2677.06 | 0.34 | 1.23e-03 |

Table S36 **Analysis of variance from the LMM of NF learning score of the control group**. We computed type III Analysis of Variance on the LMM of the Table S35 with Satterthwaite's method, using the *anova()* function of the *lmerTest* package of R.

| Parameter | Sum Squares | NumDF | DenDF | Mean Square | F | p | Eta2 (partial) |
|---|---|---|---|---|---|---|---|
| session | 404.35 | 1 | 251.03 | 404.35 | 0.51 | 0.476 | 2.02e-03 |

Additional individual linear regressions of the NF learning score (see Supplementary Fig. S8 and S9) showed that 80% (20/25) of the participants from the NF group had a positive regression slope across the 12 sessions, while the slope was positive for 48% (11/23) of the participants from the control group.

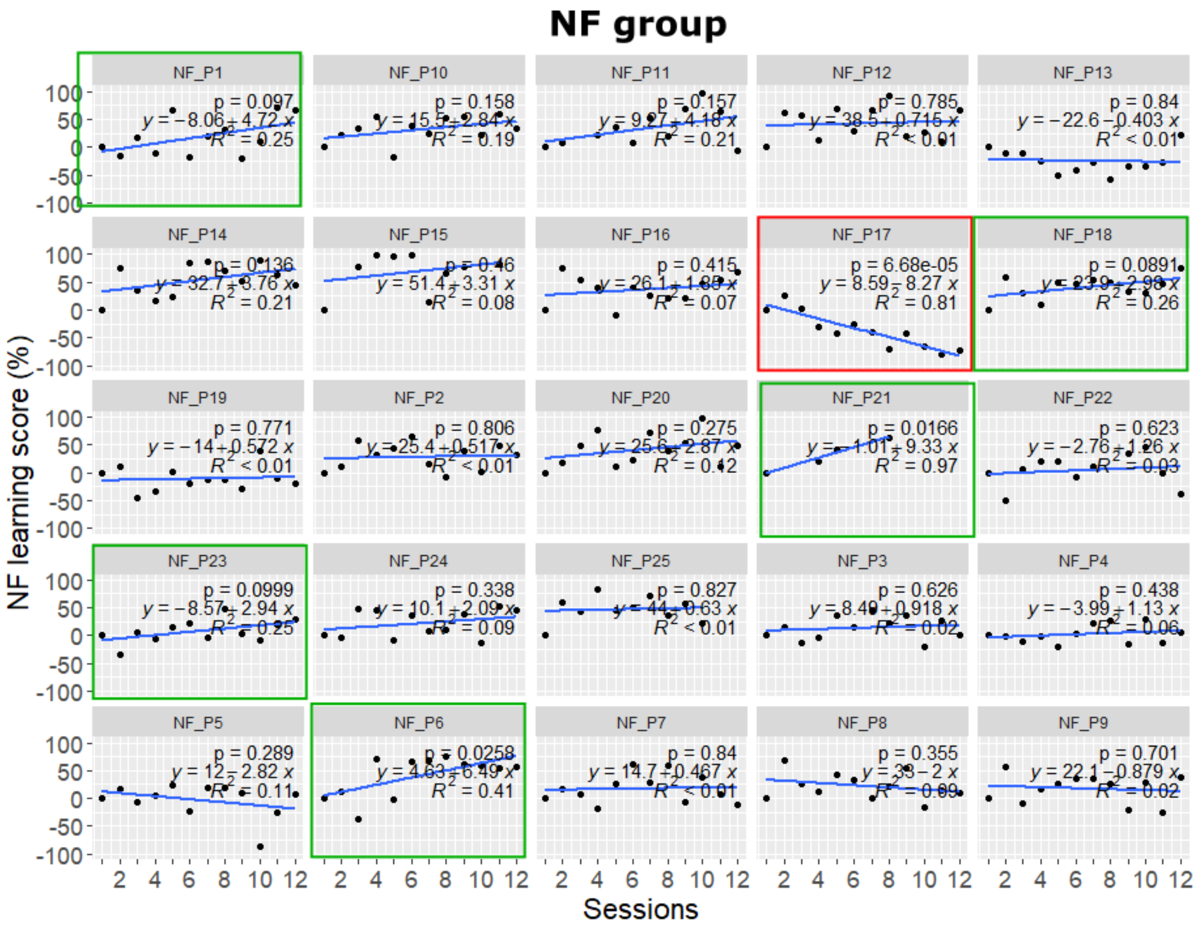


Fig. S8 **Evolution of the NF learning score across the 12 training sessions for every participant of the NF group.** Same legend as in Supplementary Fig. S6.

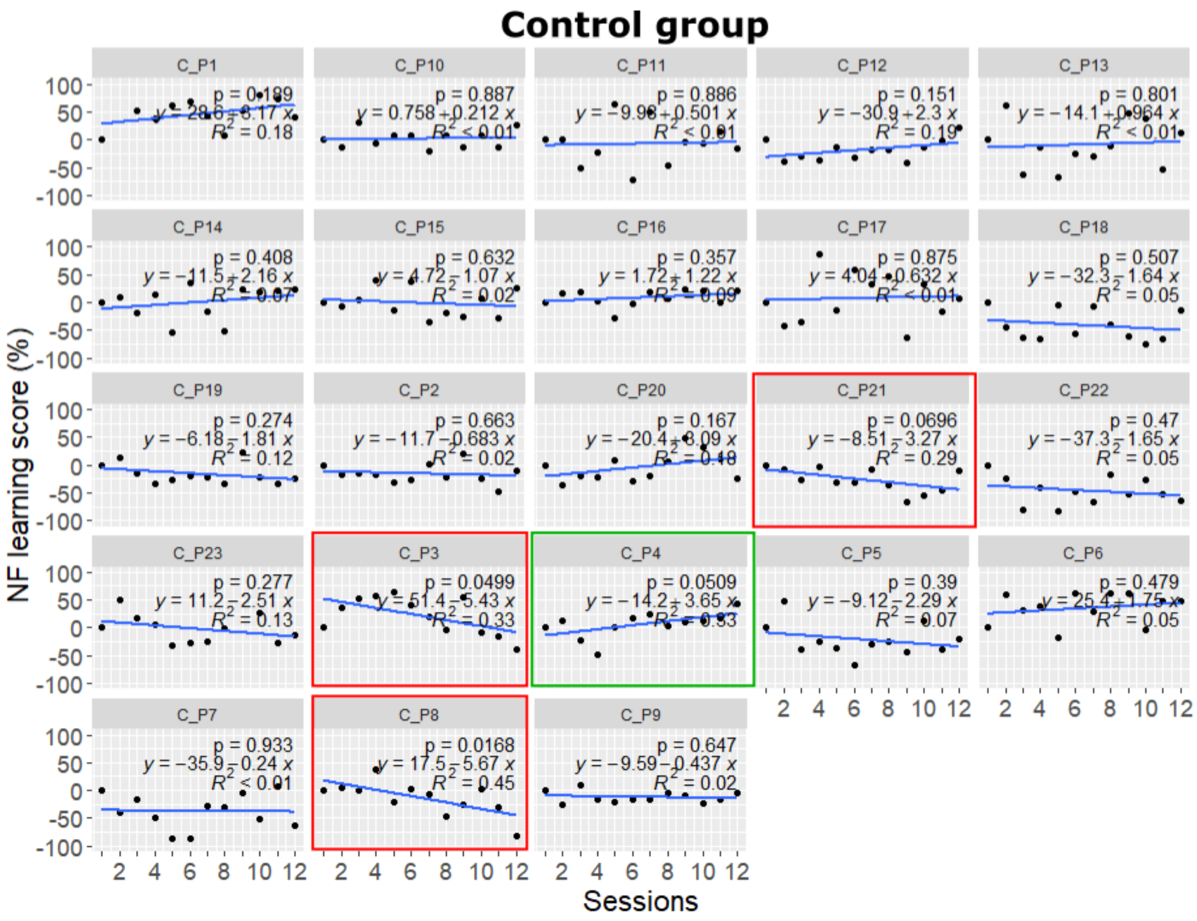


Fig. S9 **Evolution of the NF learning score across the 12 training sessions for every participant of the control group.** Same legend as in Supplementary Fig. S7.

#### 7.3. Theta and low beta band activities

##### 7.3.1. Theta band activity

Table S37 **Results of the LMM for the theta band (4-7Hz) activity analysis.** We used the LMM described in Eq. (4) of the main text with a random effects structure only composed of a random intercept. The approach was identical to that for NF learning score (described in Supplementary Table S31). See Supplementary Table S9 for table description.

| | Parameters | $\beta$ | 95% CI | |
|---|---|---|---|---|
| **Fixed effects** | (Intercept)<br>session<br>group [control]<br>session:group [control] | 4.71<br>0.06<br>0.18<br>-0.04 | [3.88, 5.54]<br>[0.02, 0.10]<br>[-1.03, 1.38]<br>[-0.10, 0.01] | |
| **Random effects** | **Parameters** | **Variance** | **Std. Dev.** | **Correlation** |
| | subject_id (intercept)<br>residual | 4.066<br>1.466 | 2.016<br>1.211 | -<br>- |
| **Fit** | **AIC** | **BIC** | **Conditional R2** | **Marginal R2** |
| | 2034.25 | 2060.36 | 0.74 | 4.25e-03 |

Table S38 **Analysis of variance from the LMM of theta band activity**. We computed type III Analysis of Variance on the LMM of the Table S37 with Satterthwaite's method, using the *anova()* function of the *lmerTest* package of R.

| Parameter | Sum Squares | NumDF | DenDF | Mean Square | F | p | Eta2 (partial) |
|---|---|---|---|---|---|---|---|
| session | 9.52 | 1 | 523.07 | 9.52 | 6.50 | **< 0.011** | 0.01 |
| group | 0.12 | 1 | 53.06 | 0.12 | 0.08 | 0.775 | 1.55e-03 |
| session:group | 3.10 | 1 | 523.07 | 3.10 | 2.11 | 0.147 | 4.02e-03 |

##### 7.3.2. Low beta band activity

Table S39 **Results of the LMM for the low beta band (13-18Hz) activity analysis.** We used the LMM described in Eq. (4) of the main text with a random effects structure only composed of a random intercept. The approach was identical to that for NF learning score (described in Supplementary Table S31). See Supplementary Table S9 for table description.

| | **Parameters** | $\beta$ | **95% CI** | |
|---|---|---|---|---|
| **Fixed effects** | (Intercept)<br>session<br>group [control]<br>session:group [control] | 3.56<br>0.01<br>0.19<br>-0.03 | [3.14, 3.97]<br>[-0.01, 0.03]<br>[-0.40, 0.79]<br>[-0.05, 0.00] | |
| **Random effects** | **Parameters** | **Variance** | **Std. Dev.** | **Correlation** |
| | subject_id (intercept)<br>residual | 1.0146<br>0.3265 | 1.0073<br>0.5714 | -<br>- |
| **Fit** | **AIC** | **BIC** | **Conditional R2** | **Marginal R2** |
| | 1184.86 | 1210.97 | 0.76 | 2.01e-03 |

Table S40 **Analysis of variance from the LMM of low beta band activity**. We computed type III Analysis of Variance on the LMM of the Table S39 with Satterthwaite's method, using the *anova()* function of the *lmerTest* package of R.

| **Parameter** | **Sum Squares** | **NumDF** | **DenDF** | **Mean Square** | **F** | **p** | **Eta2 (partial)** |
|---|---|---|---|---|---|---|---|
| session | 0.05 | 1 | 523.04 | 0.05 | 0.15 | 0.694 | 2.95e-04 |
| group | 0.13 | 1 | 52.28 | 0.13 | 0.40 | 0.531 | 7.57e-03 |
| session:group | 1.21 | 1 | 523.04 | 1.21 | 3.70 | 0.055 | 7.03e-03 |

Table S40 shows a marginal interaction between group and session. Then, to examine the effect of the session within each group, we ran additional LMM in each group separately.

Table S41 **Results of the Linear Mixed Model (LMM) for beta low band activity analysis of the NF group.** We used an LMM including a random effect structure with random intercept by participant and fixed effects for exercise and session. The model was fit using the Restricted Maximum Likelihood (REML) approach. See Supplementary Table S9 for table description.

| **Fixed effects** | **Parameters** | $\beta$ | **95% CI** | |
|---|---|---|---|---|
| | (Intercept)<br>session | 3.55<br>0.01 | [ 3.10, 4.01]<br>[-0.01, 0.03] | |
| **Random effects** | **Parameters** | **Variance** | **Std. Dev.** | **Correlation** |
| | subject_id (intercept)<br>residual | 1.2758<br>0.2861 | 1.1295<br>0.5349 | -<br>- |
| **Fit** | **AIC** | **BIC** | **Conditional R2** | **Marginal R2** |
| | 587.24 | 602.03 | 0.82 | 8.62e-04 |

Table S42 **Analysis of variance from the LMM of beta low band activity of the NF group**. We computed type III Analysis of Variance on the LMM of the Table S41 with Satterthwaite's method, using the *anova()* function of the *lmerTest* package of R.

| **Parameter** | **Sum Squares** | **NumDF** | **DenDF** | **Mean Square** | **F** | **p** | **Eta2 (partial)** |
|---|---|---|---|---|---|---|---|
| session | 0.40 | 1 | 272.03 | 0.40 | 1.40 | 0.239 | 5.10e-03 |

Table S43 **Results of the Linear Mixed Model (LMM) for the beta low band activity analysis of the control group.** We used the same LMM as for Table S41 but the data were those of the control group. The model was fit using the Restricted Maximum Likelihood (REML) approach. See Supplementary Table S9 for table description.

| | | | | |
|---|---|---|---|---|
| **Fixed effects** | **Parameters** | $\beta$ | **95% CI** | |
| | (Intercept)<br>session | 3.75<br>-0.02 | [ 3.37, 4.12]<br>[-0.04, 0.00] | |
| **Random effects** | **Parameters** | **Variance** | **Std. Dev.** | **Correlation** |
| | subject_id (intercept)<br>residual | 0.7300<br>0.3703 | 0.8544<br>0.6085 | -<br>- |
| **Fit** | **AIC** | **BIC** | **Conditional R2** | **Marginal R2** |
| | 595.66 | 610.13 | 0.66 | 2.80e-03 |

Table S44 **Analysis of variance from the LMM of the beta low band activity of the control group**. We computed type III Analysis of Variance on the LMM of the Table S43 with Satterthwaite's method, using the *anova()* function of the *lmerTest* package of R.

| Parameter | Sum Squares | NumDF | DenDF | Mean Square | F | p | Eta2 (partial) |
|---|---|---|---|---|---|---|---|
| session | 0.85 | 1 | 205.01 | 0.85 | 2.29 | 0.132 | 9.03e-03 |

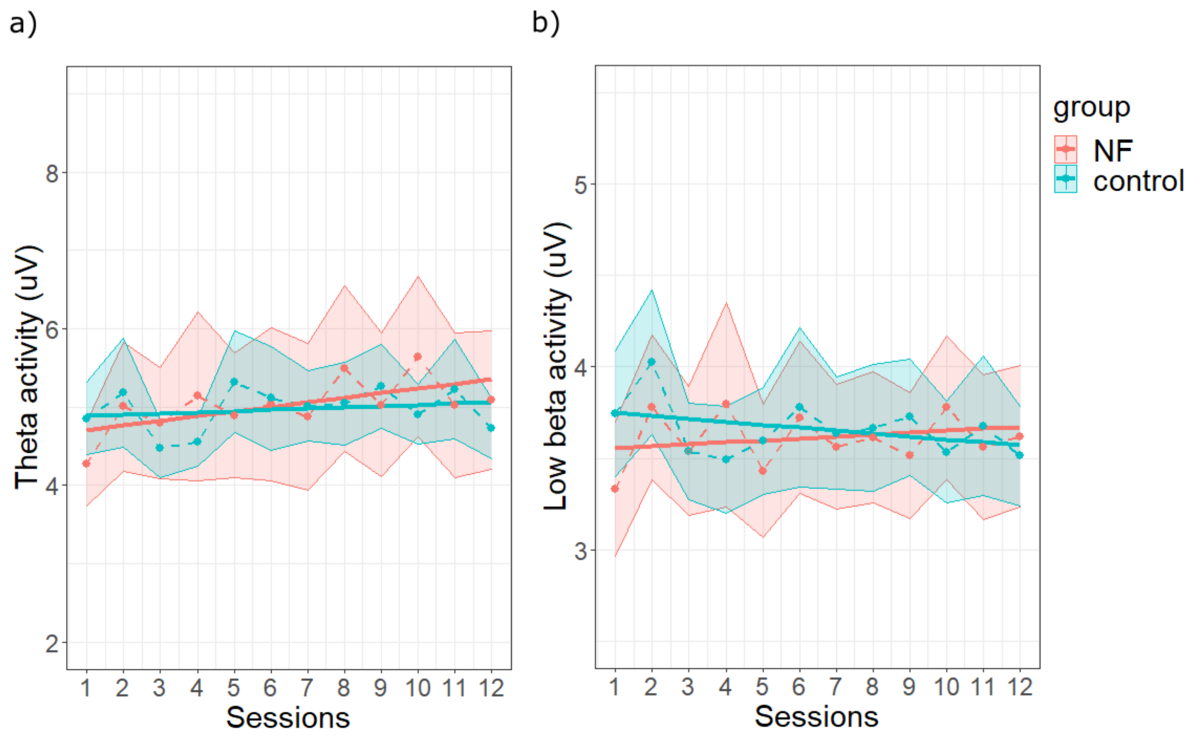


Fig. S10 **Evolution of a) theta and b) low beta band activities across sessions in the NF (in red) and the control (in blue) group.** See Fig. 3 in the main text for legend.

### 8. Quality index analysis

Table S45 **Results of the LMM for the quality index Q analysis.** We used the LMM described in Eq. (4) of the main text. The approach was identical to that for NF learning score (described in Supplementary Table S31). See Supplementary Table S9 for table description.

| | Parameters | $\beta$ | 95% CI | |
|---|---|---|---|---|
| **Fixed effects** | (Intercept)<br>session<br>group [control]<br>session:group [control] | 0.90<br>-2.02e-03<br>0.02<br>-1.06e-03 | [ 0.86, 0.93]<br>[-0.01, 0.00]<br>[-0.03, 0.06]<br>[-0.01, 0.00] | |
| **Random effects** | **Parameters** | **Variance** | **Std. Dev.** | **Correlation** |
| | subject_id (intercept)<br>session<br>residual | 5.977e-03<br>7.044e-05<br>1.323e-02 | 0.077314<br>0.008393<br>0.115016 | -<br>-0.42 (intercept)<br>- |
| **Fit** | **AIC** | **BIC** | **Conditional R2** | **Marginal R2** |
| | -5663.13 | -5612.76 | 0.31 | 5.18e-03 |

Table S46 **Analysis of variance from the LMM of the quality index Q**. We computed type III Analysis of Variance on the LMM of the Table S45 with Satterthwaite's method, using the *anova()* function of the *lmerTest* package of R.

| Parameter | Sum Squares | NumDF | DenDF | Mean Square | F | p | Eta2 (partial) |
|---|---|---|---|---|---|---|---|
| session | 0.05 | 1 | 46.130 | 0.05 | 3.70 | 0.061 | 0.07 |
| group | 5.46e-03 | 1 | 46.019 | 5.46e-03 | 0.41 | 0.524 | 8.89e-03 |
| session:group | 2.11e-03 | 1 | 46.130 | 2.11e-03 | 0.16 | 0.691 | 3.45e-03 |

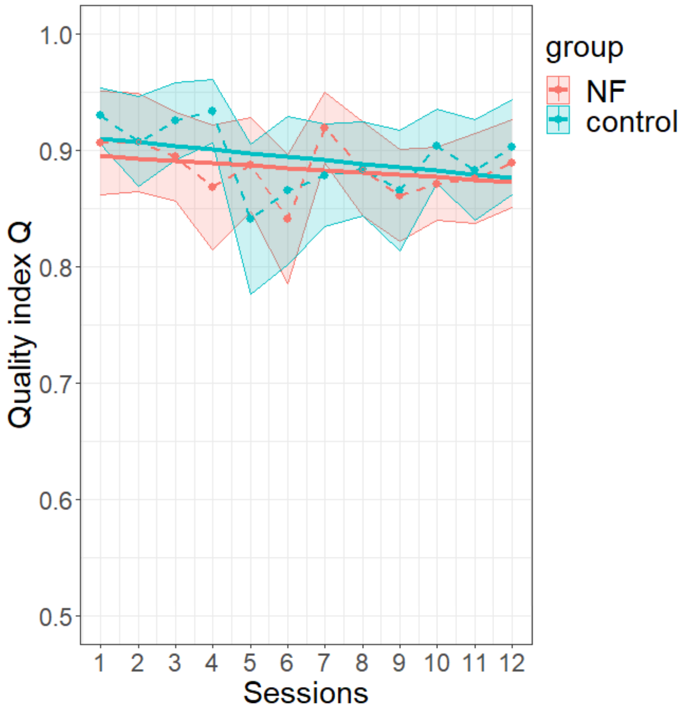


Fig. S11 **Evolution of the quality index Q across sessions in the NF (in red) and the control (in blue) groups.** See Fig. 3 in the main text for legend.

### 9. Timeline analysis

Table S47 **Results of the LMM for the timeline analysis.** We used the LMM described in Eq. (4) of the main text. The approach was identical to that for NF learning score (described in Supplementary Table S31). See Supplementary Table S9 for table description.

| | Parameters | $\beta$ | 95% CI | |
|---|---|---|---|---|
| **Fixed effects** | (Intercept)<br>session<br>group [control]<br>session:group [control] | 14.16<br>-0.05<br>0.44<br>0.07 | [12.95, 15.37]<br>[-0.12, 0.02]<br>[-1.31, 2.19]<br>[-0.04, 0.17] | |
| **Random effects** | **Parameters** | **Variance** | **Std. Dev.** | **Correlation** |
| | subject_id (intercept)<br>residual | 8.126<br>4.776 | 2.851<br>2.185 | -<br>- |
| **Fit** | **AIC** | **BIC** | **Conditional R2** | **Marginal R2** |
| | 2689.23 | 2715.35 | 0.63 | 0.01 |

Table S48 **Analysis of variance from the LMM of the timeline**. We computed type III Analysis of Variance on the LMM of the Table S47 with Satterthwaite's method, using the *anova()* function of the *lmerTest* package of R.

| Parameter | Sum Squares | NumDF | DenDF | Mean Square | F | p | Eta2 (partial) |
|---|---|---|---|---|---|---|---|
| session | 1.48 | 1 | 524.08 | 1.48 | 0.31 | 0.579 | 5.89e-04 |
| group | 1.16 | 1 | 57.50 | 1.16 | 0.24 | 0.624 | 4.20e-03 |
| session:group | 7.32 | 1 | 524.08 | 7.32 | 1.53 | 0.216 | 2.92e-03 |

### 10. Self-report outcome analysis

#### 10.1. Subjective feeling of control

Table S49 **Results of the LMM for the feeling of control analysis.** We used the LMM described in Eq. (3) of the main text with 1+session|subject_id as random effects structure. The approach was identical to that for NF index analysis. See Supplementary Table S9 for table description.

| | Parameters | $\beta$ | 95% CI | |
|---|---|---|---|---|
| **Fixed effects** | (Intercept)<br>exercise<br>session<br>group [control]<br>exercise:group [control]<br>session:group [control] | 4.16<br>0.05<br>0.13<br>-0.09<br>-0.01<br>-0.06 | [ 3.36, 4.96]<br>[ 0.02, 0.07]<br>[ 0.06, 0.19]<br>[-1.25, 1.06]<br>[-0.05, 0.02]<br>[-0.16, 0.03] | |
| | **Parameters** | **Variance** | **Std. Dev.** | **Correlation** |
| **Random effects** | subject_id (intercept)<br>session<br>residual | 3.8976<br>0.02364<br>4.06498 | 1.9742<br>0.1538<br>2.0162 | -<br>-0.34 (intercept)<br>- |
| **Fit** | **AIC** | **BIC** | **Conditional R2** | **Marginal R2** |
| | 17308.34 | 17371.28 | 0.49 | 0.02 |

Table S50 **Analysis of variance from the LMM of the feeling of control.** We computed type III Analysis of Variance on the LMM of the Table S49 with Satterthwaite's method, using the *anova()* function of the *lmerTest* package of R.

| Parameter | Sum Squares | NumDF | DenDF | Mean Square | F | p | Eta2 (partial) |
|---|---|---|---|---|---|---|---|
| exercise | 74.74 | 1 | 3905.3 | 74.74 | 18.39 | **< .001** | 4.69e-03 |
| session | 62.59 | 1 | 46.1 | 62.59 | 15.40 | **< .001** | 0.25 |
| group | 0.10 | 1 | 47.4 | 0.10 | 0.03 | 0.874 | 5.36e-04 |
| exercise:group | 2.07 | 1 | 3905.3 | 2.07 | 0.51 | 0.475 | 1.31e-04 |
| session:group | 6.59 | 1 | 46.1 | 6.59 | 1.62 | 0.209 | 0.03 |

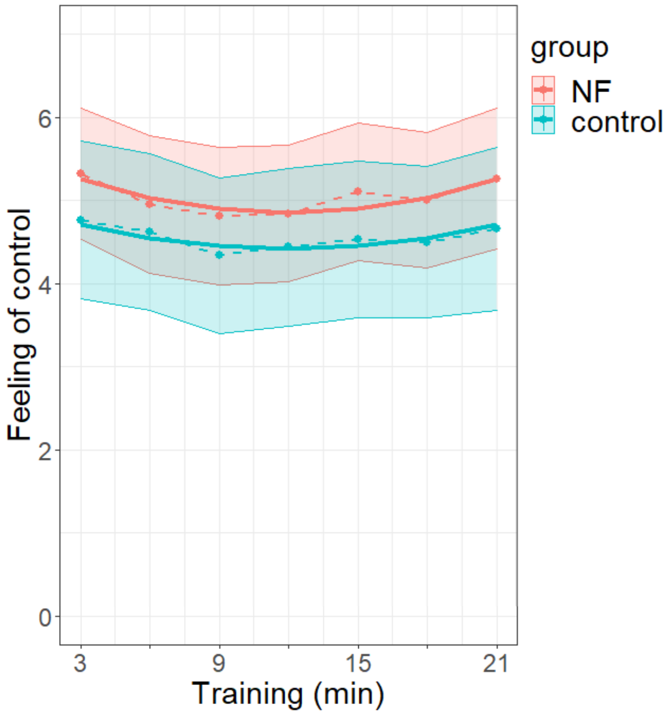


Fig. S12 **Within-session effect on the feeling of control in the NF (in red) and the control (in blue) groups.** See Fig. 4 in the main text for legend.

#### 10.2. *STAI-Y-A*

Table S51 **Results of the LMM for the *STAI-Y-A* analysis.** We used the LMM described in Eq. (5) of the main text. The approach was identical to that for NF index analysis (described in Table S25), except that the fixed effects included phase (pre-/post-session, with pre-session set as the reference phase), session, group and the two-way interactions between phase and group and between session and group. Phase [POST] denoted the overall effect of phase estimated in terms of the post- versus pre-session phase difference, phase [POST]:group [control] denoted the interaction between phase and group estimated as the difference for the control relative to the NF group in the parameter estimates of the post- versus pre-session phase effect. See Supplementary Table S9 for additional table description.

| | Parameters | $\beta$ | 95% CI | |
|---|---|---|---|---|
| **Fixed effects** | (Intercept)<br>session<br>phase [POST]<br>group [control]<br>session:group [control]<br>phase [POST]:group [control] | 32.50<br>-0.20<br>-2.48<br>1.80<br>7.36e-03<br>-2.09 | [29.16, 35.83]<br>[-0.47, 0.08]<br>[-4.41, -0.55]<br>[-3.01, 6.62]<br>[-0.39, 0.41]<br>[-4.87, 0.69] | |
| | **Parameters** | **Variance** | **Std. Dev.** | **Correlation** |
| **Random effects** | subject_id (intercept)<br>session<br>phase<br><br>residual | 67.0569<br>0.3964<br>19.3324<br><br>27.5924 | 8.1888<br>0.6296<br>4.3969<br><br>5.2528 | -<br>-0.21 (intercept)<br>[-0.60 (intercept),<br>0.30 (session)]<br>- |
| **Fit** | **AIC** | **BIC** | **Conditional R2** | **Marginal R2** |
| | 7325.55 | 7390.98 | 0.71 | 0.04 |

Table S52 **Analysis of variance from the LMM of the *STAI-Y-A*.** We computed type III Analysis of Variance on the LMM of the Table S51 with Satterthwaite's method, using the *anova()* function of the *lmerTest* package of R.

| Parameter | Sum Squares | NumDF | DenDF | Mean Square | F | p | Eta2 (partial) |
|---|---|---|---|---|---|---|---|
| session | 98.74 | 1 | 45.787 | 98.74 | 3.58 | 0.065 | 0.07 |
| phase | 683.57 | 1 | 46.137 | 683.57 | 24.77 | **< .001** | 0.35 |
| group | 3.49 | 1 | 46.017 | 3.49 | 0.13 | 0.724 | 2.74e-03 |
| session:group | 0.04 | 1 | 45.787 | 0.04 | 1.31e-03 | 0.971 | 2.86e-05 |
| phase:group | 60.17 | 1 | 46.137 | 60.17 | 2.18 | 0.147 | 0.05 |

##### 10.3. *relax-VAS*

Table S53 **Results of the LMM for the *relax-VAS* analysis.** We used the LMM described in Eq. (5) of the main text with 1+phase|subject_id as random effects structure. The approach was identical to that for *STAI-Y-A* analysis. See Supplementary Table S9 for table description.

| | Parameters | $\beta$ | 95% CI | |
|---|---|---|---|---|
| **Fixed effects** | (Intercept)<br>session<br>phase [POST]<br>group [control]<br>session:group [control]<br>phase [POST]:group [control] | 6.10<br>0.06<br>0.84<br>0.04<br>-0.02<br>0.33 | [ 5.38, 6.82]<br>[ 0.03, 0.10]<br>[ 0.37, 1.31]<br>[-1.00, 1.08]<br>[-0.07, 0.03]<br>[-0.34, 1.01] | |
| **Random effects** | **Parameters** | **Variance** | **Std. Dev.** | **Correlation** |
| | subject_id (intercept)<br>phase<br>residual | 2.986<br>1.077<br>2.055 | 1.728<br>1.038<br>1.434 | -<br>-0.66<br>- |
| **Fit** | **AIC** | **BIC** | **Conditional R2** | **Marginal R2** |
| | 4330.8 | 4381.35 | 0.56 | 0.06 |

Table S54 **Analysis of variance from the LMM of the *relax-VAS*.** We computed type III Analysis of Variance on the LMM of the Table S53 with Satterthwaite's method, using the *anova()* function of the *lmerTest* package of R.

| Parameter | Sum Squares | NumDF | DenDF | Mean Square | F | p | Eta2 (partial) |
|---|---|---|---|---|---|---|---|
| session | 38.13 | 1 | 1050.22 | 38.13 | 18.55 | **< .001** | 0.02 |
| phase | 70.48 | 1 | 46.01 | 70.48 | 34.29 | **< .001** | 0.43 |
| group | 0.42 | 1 | 55.69 | 0.42 | 0.20 | 0.653 | 3.66e-03 |
| session:group | 1.34 | 1 | 1050.22 | 1.34 | 0.65 | 0.420 | 6.20e-04 |
| phase:group | 1.91 | 1 | 46.01 | 1.91 | 0.93 | 0.340 | 0.02 |

#### 10.4. *STAI-Y-B*

Table S55 **Results of the LMM for the *STAI-Y-B* analysis.** We used an LMM with phase, group and the interaction phase:group as fixed effects and 1|subject_id as random effects structure. See Supplementary Table S9 for table description.

| | | | | |
|---|---|---|---|---|
| **Fixed effects** | **Parameters** | $\beta$ | **95% CI** | |
| | (Intercept)<br>phase [POST]<br>group [control]<br>phase [POST]:group [control] | 39.60<br>-0.85<br>1.49<br>-0.63 | [36.36, 42.84]<br>[-3.14, 1.44]<br>[-3.20, 6.17]<br>[-3.90, 2.65] | |
| **Random effects** | **Parameters** | **Variance** | **Std. Dev.** | **Correlation** |
| | subject_id (intercept)<br>residual | 52.07<br>16.42 | 7.216<br>4.053 | -<br>- |
| **Fit** | **AIC** | **BIC** | **Conditional R2** | **Marginal R2** |
| | 628.73 | 644.06 | 0.76 | 0.01 |

Table S56 **Analysis of variance from the LMM of the *STAI-Y-B*.** We computed type III Analysis of Variance on the LMM of the Table S55 with Satterthwaite's method, using the *anova()* function of the *lmerTest* package of R.

| **Parameter** | **Sum Squares** | **NumDF** | **DenDF** | **Mean Square** | **F** | **p** | **Eta2 (partial)** |
|---|---|---|---|---|---|---|---|
| phase | 31.93 | 1 | 45.273 | 31.93 | 1.94 | 0.170 | 0.04 |
| group | 4.48 | 1 | 46.039 | 4.48 | 0.27 | 0.604 | 5.89e-03 |
| phase:group | 2.32 | 1 | 45.273 | 2.32 | 0.14 | 0.709 | 3.11e-03 |

#### 10.5. *PANAS - positive*

Table S57 **Results of the LMM for the positive items of *PANAS* analysis.** The approach was identical to that for *STAI-Y-B* analysis. See Supplementary Table S9 for table description.

| | | | | |
|---|---|---|---|---|
| **Fixed effects** | **Parameters** | $\beta$ | **95% CI** | |
| | (Intercept)<br>phase [POST]<br>group [control]<br>phase [POST]:group [control] | 35.76<br>-2.18<br>-0.06<br>2.14 | [33.16, 38.36]<br>[-4.14, -0.22]<br>[-3.83, 3.70]<br>[-0.66, 4.94] | |
| **Random effects** | **Parameters** | **Variance** | **Std. Dev.** | **Correlation** |
| | subject_id (intercept)<br>residual | 32.12<br>12.03 | 5.667<br>3.469 | -<br>- |
| **Fit** | **AIC** | **BIC** | **Conditional R2** | **Marginal R2** |
| | 593.70 | 609.02 | 0.73 | 0.02 |

Table S58 **Analysis of variance from the LMM of the positive items of *PANAS* analysis.** We computed type III Analysis of Variance on the LMM of the Table S57 with Satterthwaite's method, using the *anova()* function of the *lmerTest* package of R.

| Parameter | Sum Squares | NumDF | DenDF | Mean Square | F | p | Eta2 (partial) |
|---|---|---|---|---|---|---|---|
| phase | 29.13 | 1 | 45.454 | 29.13 | 2.42 | 0.127 | 0.05 |
| group | 3.80 | 1 | 46.188 | 3.80 | 0.32 | 0.577 | 6.79e-03 |
| phase:group | 26.90 | 1 | 45.454 | 26.90 | 2.23 | 0.142 | 0.05 |

#### 10.6. *PANAS - negative*

Table S59 **Results of the LMM for the negative items of *PANAS* analysis.** The approach was identical to that for *STAI-Y-B* analysis. See Supplementary Table S9 for table description.

| | Parameters | $\beta$ | 95% CI | |
|---|---|---|---|---|
| **Fixed effects** | (Intercept)<br>phase [POST]<br>group [control]<br>phase [POST]:group [control] | 18.36<br>-0.02<br>0.12<br>-0.29 | [15.83, 20.89]<br>[-2.56, 2.53]<br>[-3.54, 3.77]<br>[-3.93, 3.35] | |
| | **Parameters** | **Variance** | **Std. Dev.** | **Correlation** |
| **Random effects** | subject_id (intercept)<br>residual | 21.26<br>20.39 | 4.611<br>4.515 | -<br>- |
| | **AIC** | **BIC** | **Conditional R2** | **Marginal R2** |
| **Fit** | 608.68 | 624.01 | 0.51 | 2.77e-04 |

Table S60 **Analysis of variance from the LMM of the negative items of *PANAS* analysis.** We computed type III Analysis of Variance on the LMM of the Table S59 with Satterthwaite's method, using the *anova()* function of the *lmerTest* package of R.

| Parameter | Sum Squares | NumDF | DenDF | Mean Square | F | p | Eta2 (partial) |
|---|---|---|---|---|---|---|---|
| phase | 0.60 | 1 | 45.494 | 0.60 | 0.03 | 0.864 | 6.51e-04 |
| group | 5.25e-03 | 1 | 46.012 | 5.25e-03 | 2.58e-04 | 0.987 | 5.60e-06 |
| phase:group | 0.49 | 1 | 45.494 | 0.49 | 0.02 | 0.877 | 5.30e-04 |

#### 10.7. *PSS*

Table S61 **Results of the LMM for the *PSS* analysis.** The approach was identical to that for *STAI-Y-B* analysis. See Supplementary Table S9 for table description.

| | Parameters | $\beta$ | 95% CI | |
|---|---|---|---|---|
| **Fixed effects** | (Intercept)<br>phase [POST]<br>group [control]<br>phase [POST]:group [control] | 37.08<br>-0.24<br>0.49<br>-1.46 | [34.25, 39.91]<br>[-2.57, 2.10]<br>[-3.60, 4.57]<br>[-4.80, 1.88] | |
| | **Parameters** | **Variance** | **Std. Dev.** | **Correlation** |
| **Random effects** | subject_id (intercept)<br>residual | 34.96<br>17.11 | 5.913<br>4.137 | -<br>- |
| **Fit** | **AIC** | **BIC** | **Conditional R2** | **Marginal R2** |
| | 615.65 | 630.97 | 0.67 | 7.14e-03 |

Table S62 **Analysis of variance from the LMM of the *PSS* analysis.** We computed type III Analysis of Variance on the LMM of the Table S61 with Satterthwaite's method, using the *anova()* function of the *lmerTest* package of R.

| Parameter | Sum Squares | NumDF | DenDF | Mean Square | F | p | Eta2 (partial) |
|---|---|---|---|---|---|---|---|
| phase | 21.99 | 1 | 45.270 | 21.99 | 1.29 | 0.263 | 0.03 |
| group | 0.28 | 1 | 45.948 | 0.28 | 0.02 | 0.899 | 3.57e-04 |
| phase:group | 12.55 | 1 | 45.270 | 12.55 | 0.73 | 0.396 | 0.02 |

### 11. Group comparison at the first session

Based on Fig. 3, one could suspect a difference between groups at the first session for the NF index values. We first plotted the individual progressions in each group in order to visually assess if the significant difference between groups in the progression of NF index across sessions seemed due to lower values of the NF index at the first session in the NF versus the control group. For this, we displayed, for each user, the linear regression fits of NF index across the sessions with a color that coded the observed mean value of the NF index at the first session (S1):

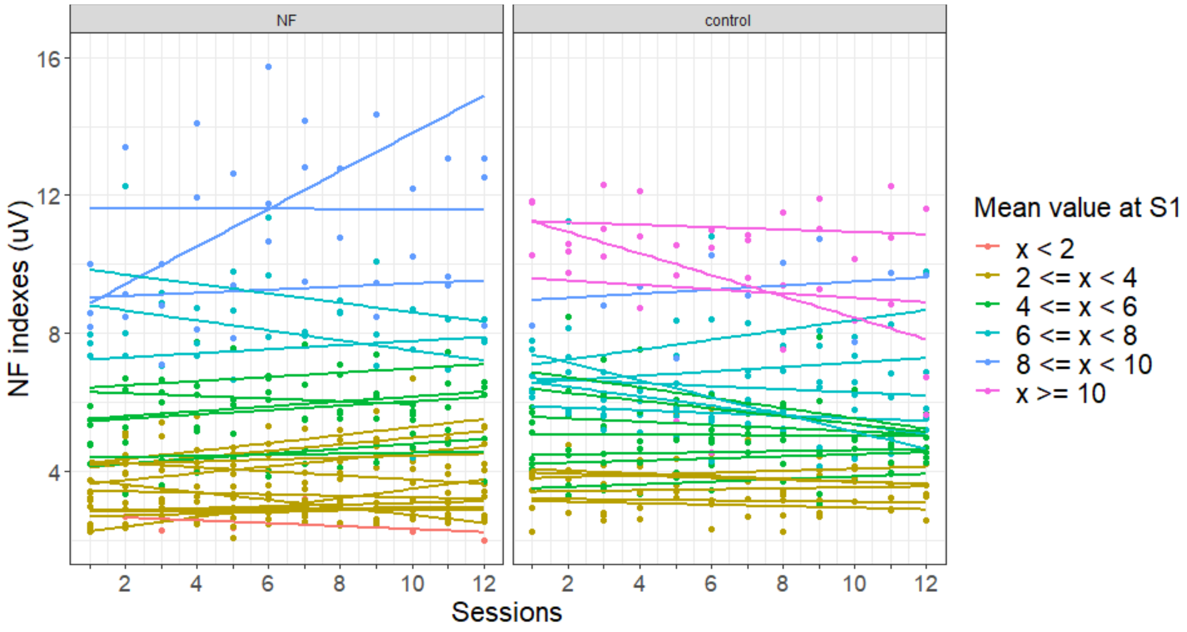


Fig. S13: **Individual linear fits of NF index values across sessions according to the mean value of NF index at the first session.** Points are the mean NF index value of each subject at each session.

Fig. S13 indicated no clear difference between groups at the first session and the starting NF index value did not seem to systematically influence the progression of the NF index across sessions.

To test statistically the variability between groups at the first session, we performed, for each outcome variable of interest, an independent t-test between groups (Table S63). All these statistical tests were performed with the *t_test()* function of the *rstatix* package of R. Note that we did not correct for multiple comparisons here in order to set a non-conservative threshold for detecting such potential initial difference.

Table S63 **Independent t-tests between groups at the first session**. Std. Dev. is the standard deviation; *n* is the number of subjects; *p* is the p-value obtained with the t-test (t); df is the degrees of freedom; d is the Cohen's distance to assess the effect size.

| Outcome variable | Groups | mean | Std. Dev. | n | t | df | p | d |
|---|---|---|---|---|---|---|---|---|
| **NF index** | NF | 4.71 | 2.34 | 48 | -1.8 | 46 | 0.0789 | -0.519 |
| | control | 6 | 2.62 | | | | | |
| **Theta activity** | NF | 4.27 | 1.69 | 48 | -1.27 | 46 | 0.212 | -0.366 |
| | control | 4.85 | 1.47 | | | | | |
| **Low beta activity** | NF | 3.33 | 1.17 | 48 | -1.27 | 46 | 0.212 | -0.366 |
| | control | 3.74 | 1.09 | | | | | |
| **STAI-Y-A** | NF | 32.2 | 7.65 | 48 | -0.545 | 46 | 0.588 | -0.158 |
| | control | 33.5 | 8.6 | | | | | |
| **relax-VAS** | NF | 6.64 | 2.07 | 48 | 0.672 | 46 | 0.505 | 0.194 |
| | control | 6.2 | 2.4 | | | | | |
| **Quality index** | NF | 0.906 | 0.124 | 48 | -0.828 | 46 | 0.412 | -0.239 |
| | control | 0.93 | 0.058 | | | | | |
| **Feeling of control** | NF | 4.03 | 2.37 | 48 | 0.956 | 46 | 0.344 | 0.276 |
| | control | 3.38 | 2.32 | | | | | |
| **STAI-Y-B** | NF | 39.6 | 8.18 | 48 | -0.590 | 46 | 0.588 | -0.170 |
| | control | 41.1 | 9.28 | | | | | |
| **PANAS-positive** | NF | 35.8 | 5.73 | 48 | 0.0382 | 46 | 0.97 | 0.011 |
| | control | 35.7 | 5.94 | | | | | |
| **PANAS-negative** | NF | 18.4 | 5.98 | 48 | -0.0671 | 46 | 0.947 | -0.0194 |
| | control | 18.5 | 6.24 | | | | | |
| **PSS** | NF | 37.1 | 6.47 | 48 | -0.248 | 46 | 0.805 | -0.0718 |
| | control | 37.6 | 7.07 | | | | | |

Table S63 shows that there was not any significant difference between groups at the first session, for either studied variable (all uncorrected *p>0.05*).

### 12. Study of LOWq, MEDq and HIGHq proportions

To study the evolution of the LOWq, MEDq and HIGHq EEG segments between the first and last sessions, we did the cumulative sum of the number of segments in each category across both channels and all the subjects. Then, a proportion was obtained by dividing this cumulative sum by the total number of EEG segments for the first and the last session respectively (see Supplementary Table S64).

Table S64 **Proportion of LOWq, MEDq and HIGHq EEG segments in the first and last sessions across both channels and all the subjects**

| | **LOWq** | **MEDq** | **HIGHq** |
|---|---|---|---|
| **First session** | 0.0127 | 0.1229 | 0.8643 |
| **Last session** | 0.0187 | 0.1479 | 0.8333 |

A chi-squared goodness of fit analysis was performed with the *chisq.test()* function of the *stats* package of R to analyze the evolution of these proportions between the first and the last sessions. No significant change was observed ($X^2$ = 0.0090422, df = 2, $p$ = 0.9955).

#### 13. Correlations between NF index and self-report outcomes

For each participant, an NF index averaged value across exercises was obtained for each session. Similarly, the averaged value for each session was obtained for the feeling of control score. Concerning *STAI-Y-A* and *relax-VAS* scores, pre- and post-session scores were averaged for each participant on each session. Pearson's correlation coefficients were calculated between NF index averaged values and self-report outcome averaged values (*STAI-Y-A*, *relax-VAS* and feeling of control) for each session and each group.

In addition, for each participant, the slopes of the linear regressions across sessions were also computed from the above described averaged values, for NF index and self-report outcomes. Pearson's correlation coefficients were computed between the slopes of NF index outcome and the slopes of the self-report outcomes, in each group.

To adjust for multiple comparisons, an approximate multivariate permutation test was conducted. Sampling distribution was built to calculate the corrected p value as the proportion of values that were larger than the observed correlation coefficient value (Nichols et Holmes, 2002).

Table S65 **Pearson’s correlation analyses between NF index outcome and *relax-VAS* outcome for each group.** The first 12 rows correspond to correlations between averaged values at each session and the 13th row corresponds to the correlation between the slopes. *r* is Pearson's correlation coefficient; *p_val* is the p-value obtained from Pearson's correlation analyses and *p_corr* is the corrected p-value after multiple comparisons adjustment.

| | ***NF group*** | | | ***control group*** | | |
|---|---|---|---|---|---|---|
| | ***r*** | ***p_val*** | ***p_corr*** | ***r*** | ***p_val*** | ***p_corr*** |
| **S1** | -0.399 | **0.031** | 0.272 | 0.01 | 0.481 | 0.944 |
| **S2** | -0.028 | 0.445 | 1 | 0.233 | 0.145 | 0.86 |
| **S3** | -0.167 | 0.211 | 1 | 0.064 | 0.383 | 0.936 |
| **S4** | -0.298 | 0.078 | 0.592 | 0.066 | 0.38 | 0.912 |
| **S5** | -0.132 | 0.26 | 0.978 | 0.02 | 0.462 | 0.972 |
| **S6** | -0.092 | 0.326 | 0.986 | -0.295 | 0.093 | 0.756 |
| **S7** | -0.055 | 0.393 | 1 | -0.061 | 0.389 | 1 |
| **S8** | -0.022 | 0.457 | 1 | -0.17 | 0.219 | 1 |
| **S9** | 0.031 | 0.439 | 0.942 | 0.229 | 0.147 | 0.858 |
| **S10** | -0.12 | 0.28 | 1 | -0.188 | 0.197 | 0.982 |
| **S11** | -0.089 | 0.333 | 1 | 0.035 | 0.436 | 0.934 |
| **S12** | -0.016 | 0.467 | 1 | -0.108 | 0.306 | 1 |
| **Slope** | 0.045 | 0.412 | 0.976 | 0.107 | 0.312 | 1 |

Table S66 **Pearson's correlation analyses between NF index outcome and *STAI-Y-A* outcome for each group.** See Supplementary Table S65 for table description.

| | ***NF group*** | | | ***control group*** | | |
|---|---|---|---|---|---|---|
| | ***r*** | ***p_val*** | ***p_corr*** | ***r*** | ***p_val*** | ***p_corr*** |
| **S1** | 0.378 | **0.02** | 0.364 | -0.079 | 0.178 | 1 |
| **S2** | 0.068 | 0.185 | 0.97 | -0.125 | 0.141 | 0.998 |
| **S3** | 0.021 | 0.229 | 0.932 | -0.205 | 0.087 | 0.878 |
| **S4** | 0.15 | 0.118 | 0.996 | -0.125 | 0.141 | 0.99 |
| **S5** | 0.049 | 0.202 | 0.966 | -0.035 | 0.218 | 1 |
| **S6** | -0.039 | 0.212 | 0.988 | 0.025 | 0.227 | 0.998 |
| **S7** | -0.061 | 0.191 | 1 | -0.019 | 0.231 | 0.99 |
| **S8** | -0.083 | 0.172 | 1 | 0.103 | 0.159 | 1 |
| **S9** | -0.001 | 0.248 | 1 | -0.257 | 0.059 | 0.748 |
| **S10** | -0.165 | 0.107 | 0.96 | 0.037 | 0.215 | 1 |
| **S11** | -0.03 | 0.221 | 1 | -0.135 | 0.134 | 0.998 |
| **S12** | 0.087 | 0.168 | 1 | 0.17 | 0.107 | 0.992 |
| **Slope** | -0.069 | 0.185 | 0.94 | -0.103 | 0.158 | 0.982 |

Table S67 **Pearson's correlation analyses between NF index outcome and feeling of control for each group.** See Supplementary Table S65 for table description.

| | ***NF group*** | | | ***control group*** | | |
|---|---|---|---|---|---|---|
| | ***r*** | ***p_val*** | ***p_corr*** | ***r*** | ***p_val*** | ***p_corr*** |
| **S1** | -0.038 | 0.213 | 1 | -0.237 | 0.08 | 0.608 |
| **S2** | 0.125 | 0.135 | 0.996 | -0.117 | 0.252 | 0.944 |
| **S3** | 0.298 | **0.039** | 0.692 | -0.152 | 0.185 | 0.864 |
| **S4** | -0.02 | 0.23 | 1 | -0.239 | 0.081 | 0.624 |
| **S5** | 0.281 | **0.045** | 0.786 | -0.102 | 0.277 | 1 |
| **S6** | 0.101 | 0.157 | 0.97 | -0.113 | 0.257 | 1 |
| **S7** | -0.013 | 0.237 | 0.986 | -0.283 | 0.051 | 0.452 |
| **S8** | 0.062 | 0.191 | 0.982 | -0.113 | 0.256 | 0.994 |
| **S9** | 0.251 | 0.059 | 0.798 | -0.257 | 0.073 | 0.622 |
| **S10** | -0.083 | 0.172 | 1 | -0.289 | **0.048** | 0.424 |
| **S11** | -0.086 | 0.17 | 0.95 | -0.146 | 0.199 | 1 |
| **S12** | -0.096 | 0.16 | 0.99 | 0.038 | 0.412 | 0.996 |
| **Slope** | -0.198 | 0.086 | 1 | 0.092 | 0.292 | 1 |
